# Calibrated high-throughput electrophysiology enables clinical interpretation of *CACNA1G* missense variants

**DOI:** 10.64898/2026.05.10.724145

**Authors:** Rocio K. Finol-Urdaneta, Chek-Ying Tan, Neven Maksemous, Joanne G. Ma, Paul J. Lockhart, Penny Snell, Abhishek Malhotra, Bryony A. Thompson, Gunjan Garg, Himanshu Goel, Lyn R. Griffiths, David J. Adams, Jamie I Vandenberg, Chai-Ann Ng

## Abstract

**Objective:** Accurate classification of ion channel variants of uncertain significance (VUS) remains a persistent challenge in clinical genomics, limiting diagnostic resolution in neurological disorders.

**Methods:** We developed a calibrated electrophysiological framework to generate functional evidence for clinical interpretation of *CACNA1G* variants encoding the low-voltage-activated calcium channel Cav3.1. Functional metrics derived from automated patch⍰clamp recordings were calibrated against benign/likely benign (B/LB) and pathogenic/likely pathogenic (P/LP) reference variants to enable conservative application of ACMG/AMP functional criteria within clinical variant interpretation workflows.

**Results:** Calibration using 25 B/LB and 16 P/LP *CACNA1G* variants showed that more than 80% of P/LP variants exhibited reduced current density (CD). Deactivation kinetics (τ_Deact_) provided complementary discriminatory information by identifying gating abnormalities in variants with preserved CD. Application of this dual-metric framework to five VUS identified in Australian patients revealed two variants (Cav3.1-R186Q and R1394Q) with abnormal functional profiles consistent with voltage-sensor perturbation, supporting reassessment as likely pathogenic under ACMG/AMP guidelines. The remaining VUS displayed functional properties overlapping the benign reference distribution.

**Conclusion:** These findings establish a calibrated functional framework for generating electrophysiological evidence that supports clinical interpretation of *CACNA1G* missense variants under ACMG/AMP guidelines. When applied as external functional evidence, this approach improves resolution of *CACNA1G*-associated VUS while maintaining conservative standards for variant classification.

---

More than half of the 3.7 million variants catalogued in ClinVar are classified as variants of uncertain significance (VUS),^1^ reflecting insufficient evidence for classification under the American College of Medical Genetics and Genomics (ACMG) and the Association for Molecular Pathology (AMP) guidelines.^2^ The high prevalence of VUS represents a major challenge in genomic medicine, particularly for rare diseases such as ion channelopathies. The high prevalence of VUS represents a major challenge in genomic medicine, particularly for rare diseases such as ion-channelopathies, where diagnostic decision-making often depends on functional evidence. Among ion channel genes, *CACNA1G*, which encodes the Cav3.1 T-type calcium channel, has emerged as a gene of significant clinical interest. Pathogenic variants in *CACNA1G* are an established cause of spinocerebellar ataxia type 42 (SCA42),^3–5^ and may act as genetic modifiers in epilepsy syndromes associated with *SCN1A*-Dravet syndrome^6^ or *SCN2A.*^7^ More broadly, variants in *CACNA1G* have also been implicated in neurodevelopmental disorders.^8–10^

Cav3.1 channels are highly expressed in the cerebellum and thalamus, where they regulate rhythmic oscillations, burst firing, and neuronal excitability. These processes are essential for motor coordination, sleep, and cognition.^7, 11^ In addition, Cav3.1 activity in the medial prefrontal cortex serves as a critical driver of GABAergic neuron activation, contributing to chronic psychobiological stress responses in mice.^12^ Pathogenic *CACNA1G* variants can result in either gain- or loss-of-function effects on channel activity; however, the functional consequences of most missense variants remain unknown.^9, 13^ Of the 896 *CACNA1G* missense variants currently listed in ClinVar, 682 are classified as VUS (accessed 11 Apr 2026), underscoring the need for calibrated functional evidence to support clinical interpretation.

Interpretation of functional data in isolation can be challenging, as modest deviations from canonical “wild⍰type” behavior may overlap with normal experimental or biological variation. Anchoring electrophysiological measurements to benign reference distributions therefore provides important interpretive context. High⍰throughput automated patch⍰clamp (APC) electrophysiology offers a scalable approach for generating functional evidence relevant to ACMG/AMP⍰based variant classification.^2^ When calibrated against well⍰characterized benign and pathogenic control variants, functional data derived from APC assays can be applied at strong or moderate evidence levels under ClinGen Sequence Variant Interpretation recommendations (PS3/BS3).^14, 15^ Such calibrated APC⍰based frameworks have been applied in other clinical variant⍰interpretation settings,^14, 16–19^ but have not yet been systematically established for *CACNA1G*⍰associated neurological phenotypes.

Here, we present the first calibrated APC-based functional framework for *CACNA1G*, incorporating 25 benign and 16 pathogenic control variants to establish biologically meaningful thresholds for loss- or gain-of-function on Cav3.1 channel function. Using this framework, we evaluated five *CACNA1G* VUS identified in Australian patients with neurological phenotypes. Functional analyses revealed variant-specific effects on current density and channel gating properties. By generating calibrated functional evidence suitable for conservative application under the PS3 criterion of the ACMG/AMP guidelines, this study provides a practical approach to improve *CACNA1G* variant interpretation and reduce diagnostic uncertainty for channelopathy-associated disorders.

## Methods

### Curation of Variant Controls

Variants were curated using transcript NM_018896.5 on the GRCh38 (hg38) reference genome and classified according to American College of Medical Genetics and Genomics and Association for Molecular Pathology (ACMG/AMP) criteria ^2^. Population frequency data were obtained from gnomAD v3.1.2 (Genomes) and v2.1.1 (Exomes), excluding the Amish cohort (allele count <1,000), and were updated to gnomAD v4.1 following its release. A minor allele frequency (MAF) threshold of 0.00001 was applied, consistent with the estimated prevalence of dominant hereditary cerebellar ataxias;^20^ variants exceeding this threshold were assigned BS1. Criterion BS2 was applied at strong or supporting levels based on homozygous or heterozygous occurrence in healthy individuals. Pathogenic criteria included PS2 or PM6 for confirmed or assumed *de novo* variants (ClinGen SVI v1.1), PS4 (moderate) for variants reported in multiple unrelated affected individuals and absent from population controls, PM2 (supporting) for variants rare or absent in gnomAD, and PP3 based on concordant *in silico* predictions (REVEL, SIFT, PolyPhen-2, MutationTaster); REVEL scores were not used in isolation. Functional evidence was deliberately excluded from control variant classification to ensure independence between clinical curation and subsequent assay calibration. The final reference dataset therefore comprised 25 benign/likely benign (B/LB) and 16 pathogenic/likely pathogenic (P/LP) *CACNA1G* variant controls (Table S1).

### Cloning, Cell Line Generation and Cell Culture

Flp-In *CACNA1G* HEK293 cell lines were generated as described by Ng *et al*.^21^ *CACNA1G* variant constructs were subcloned into pcDNA™5/FRT/TO (Thermo Fisher Scientific) by Genscript Inc. (Piscataway, USA). For transfection, 60 ng *CACNA1G* plasmid DNA was co-transfected with 520 ng pOG44 Flp-Recombinase Expression Vector (Thermo Fisher Scientific, cat. #V600520) into Flp-In HEK293 cells (Thermo Fisher Scientific, cat. #R78007) using Lipofectamine 3000 (Thermo Fisher Scientific, cat. #L3000015) according to the manufacturer’s instructions. Stable integrants were selected using 200 μg/mL Hygromycin (Thermo Fisher Scientific, cat. #10687010) in DMEM (Thermo Fisher Scientific, cat. #10566024) supplemented with 10% foetal bovine serum and 10 μg/mL Blasticidin (InvivoGen, cat. #ant-bl-1). Cells were maintained at 37 °C in a humidified incubator with 5% CO₂. Prior to electrophysiological recordings, cells were harvested by enzymatic digestion following established protocols.^21^

### Functional Characterization

Electrophysiological recordings of Cav3.1-mediated calcium currents were performed using the SyncroPatch 384PE system (Nanion Technologies), equipped with low-resistance, large-hole planar patch-clamp chips optimised for reproducible, high-throughput measurements. This platform was used to generate standardized functional metrics suitable for downstream variant interpretation. Cells were voltage-clamped using solutions designed to isolate T-type calcium currents. The internal solution contained (in mM): 10⍰NaCl, 60⍰CsCl, 60⍰CsF, 10⍰HEPES, and 10⍰EGTA, adjusted to pH⍰7.2 with CsOH and an osmolarity of 285⍰±⍰3⍰mOsm. The external solution consisted of (in mM): 20⍰NaCl, 5⍰KCl, 1⍰MgCl₂, 10⍰CaCl₂, 110⍰TEA-Cl, 10⍰HEPES, and 5⍰glucose, adjusted to pH⍰7.4 with TEA-OH and an osmolarity of 295⍰±⍰3⍰mOsm.

Each experimental plate included wild⍰type (WT) and blank (negative; NEG) controls to ensure assay fidelity and enable normalization across recordings. Variants were assessed in batches of ten per run and co⍰analysed with WT and NEG controls to control for inter⍰plate variability and maintain internal consistency (see Fig 1B). Voltage⍰clamp protocols used to assess Cav3.1 channel function are shown in Figures 2, 4, S1, and⍰S2. All recordings were obtained from a holding potential of −100 mV using standardized activation, inactivation, recovery, and deactivation paradigms.

**Figure 1.**
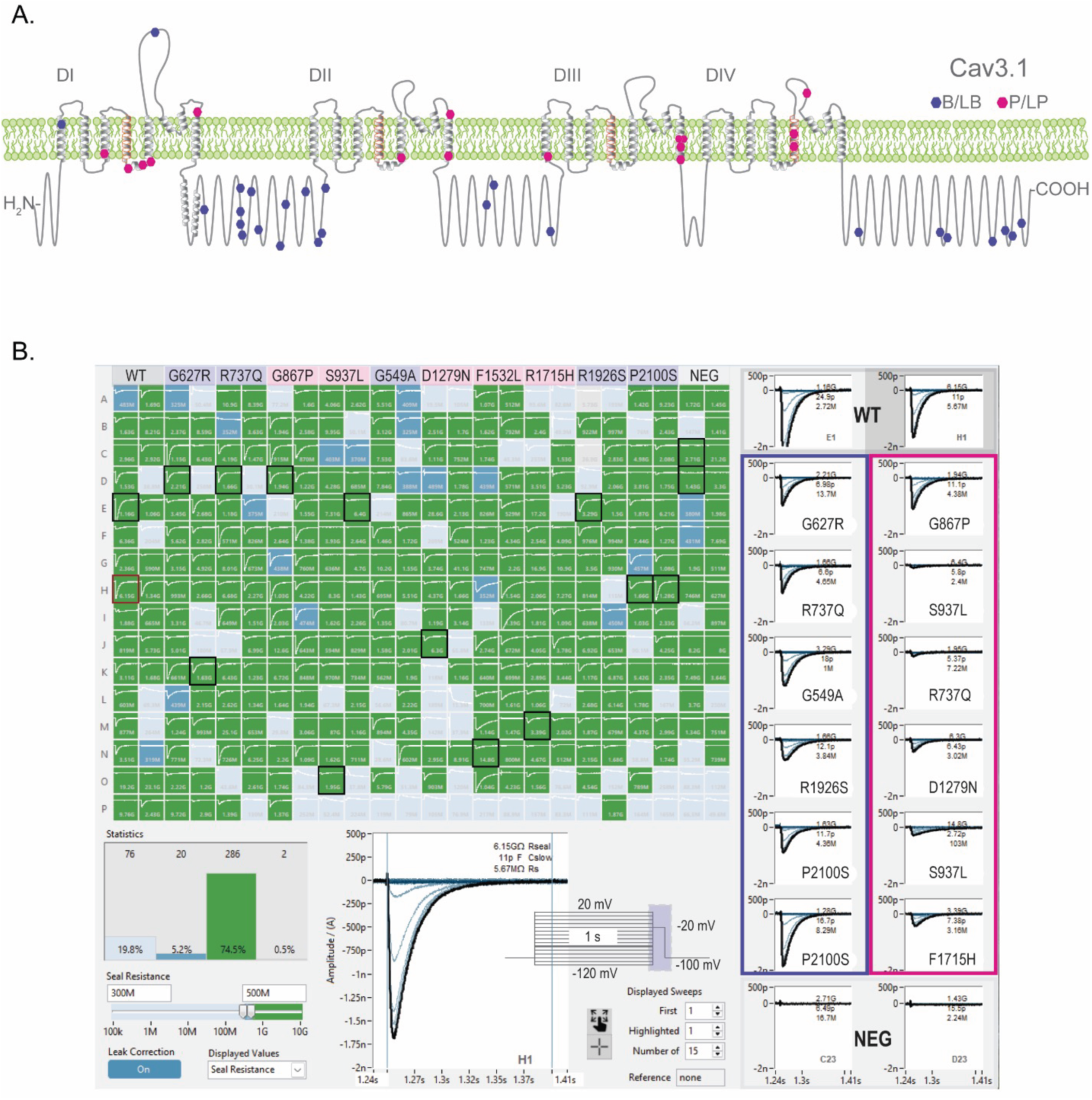
Distribution of benign and pathogenic CACNA1G amino acid substitutions and overview of the functional asay framework. **A.** Schematic representation of the Cav3.1 protein illustrating transmembrane segments (S1–S6; S4 highlighted in orange), pore loops, and cytoplasmic regions. Positions of amino⍰acid substitutions corresponding to benign/likely benign (B/LB, blue) and pathogenic/likely pathogenic (P/LP, magenta) variants are indicated to show their distribution across channel domains. **B.** Overview of the automated patch⍰clamp recording workflow used to generate standardized electrophysiological measurements. Each experimental run included Cav3.1 wild⍰type (WT)–expressing cells and blank (NEG) controls, with recording plates configured to assess 10 variants per run alongside internal controls to support calibration and normalization.

**Figure 2.**
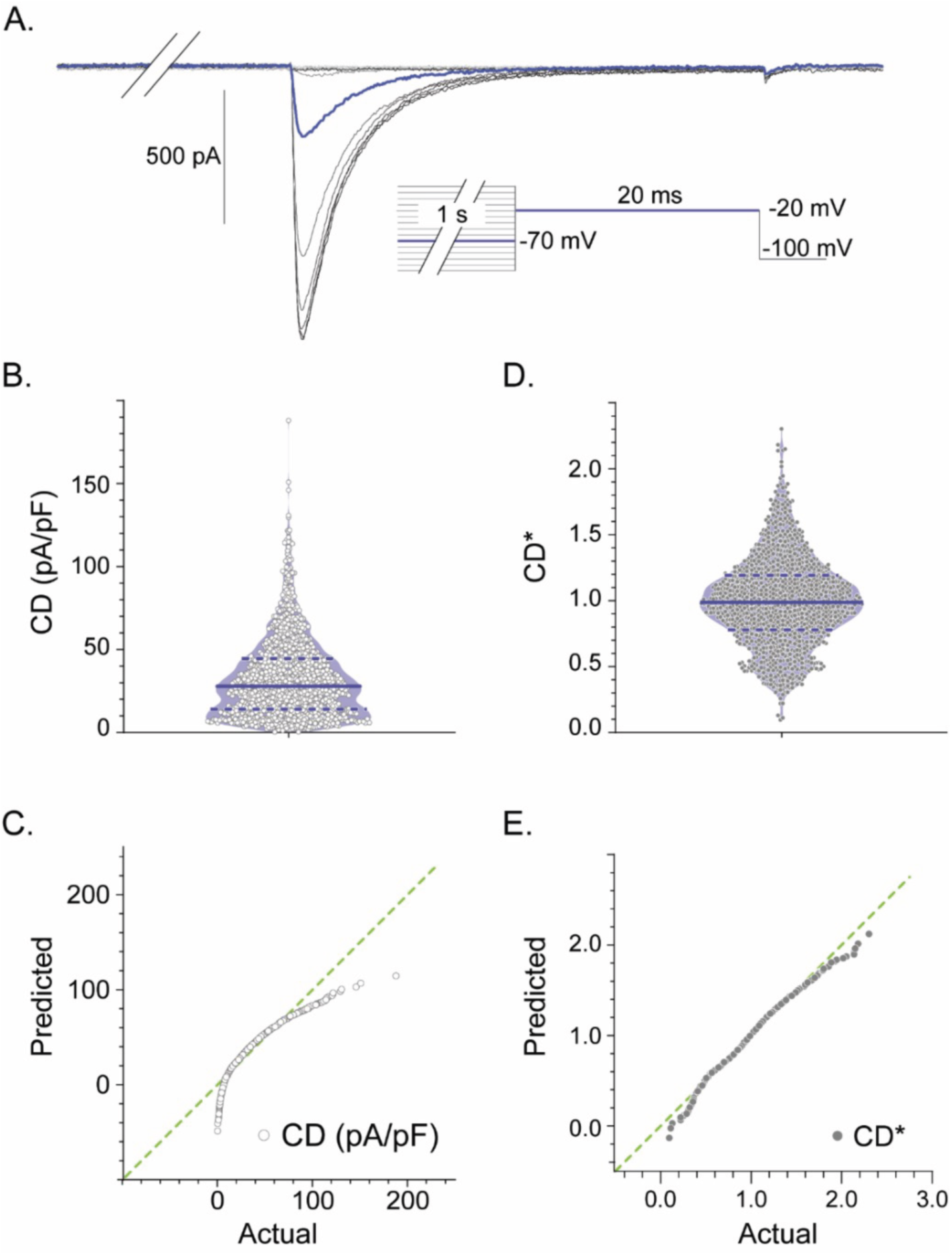
Distribution of Cav3.1-WT currents measured by automated patchlZIclamp. **A.** Representative whole-cell current traces recorded from a Flp-In™ HEK293 cell stably expressing the canonical *CACNA1G* transcript (NM_018896.5). Currents were elicited by a −20⍰mV test pulse following 1⍰s conditioning pre⍰pulses from −120⍰mV to +20⍰mV in 10⍰mV increments (protocol shown in inset). Peak inward currents measured during the −20⍰mV test pulse after a −70⍰mV pre⍰pulse (highlighted in blue) were normalized to cell capacitance to derive current density (CD). **B–C.** Violin and quantile–quantile (Q-Q) plots of raw Cav3.1-WT current density (n = 1179 cells, 62 experiments), demonstrating deviation from normality (Kolmogorov–Smirnov [KS] distance = 0.09758, p < 0.0001). **D–E.** Violin and Q-Q plots of square root–transformed current density (CD*), showing improved conformity to a Gaussian distribution (KS distance = 0.03438, p = 0.0023). Median values (solid blue line) and interquartile ranges (dotted blue lines) are indicated.

Current Density (CD) was calculated by dividing the peak inward current recorded at −20 mV by the measured cell capacitance (pA/pF). Currents were elicited following 1 s pre-pulses ranging from −120 mV to 20 mV in 10 mV increments (Fig 2A). Among these, the −70 mV pre-pulse consistently yielded the most physiologically relevant and discriminative results, approximating the resting membrane potential of central neurons and reflecting conditions under which Cav3.1 channels are likely to contribute to neuronal excitability in vivo.

Voltage dependence of activation and inactivation was assessed by plotting normalized macroscopic conductance or current amplitude against command voltage (Fig S1). Conductance was calculated as peak current divided by the driving force. Activation and steady-state inactivation curves were fitted with a Boltzmann equation (See Equations used in APC Analysis). Kinetic parameters were extracted from current traces evoked by a depolarizing step to −20⍰mV. Activation kinetics were quantified as time-to-peak (ms), measured from the onset of the voltage step to the point of maximal inward current (Fig S2). Inactivation kinetics were approximated using the relative current remaining at 50⍰ms (Irel@50ms), defined as the ratio of current at 50⍰ms to peak current (I_50ms / I_peak) (Fig S2). These surrogate measures were selected to avoid artifacts from exponential fitting, particularly for low current amplitude or traces with slow kinetics.

Deactivation kinetics (τ_Deact_) were assessed by fitting a single exponential to the decay of tail currents recorded upon repolarization to −100⍰mV following 10⍰ms depolarizing steps from −120⍰mV to 50⍰mV. Although tail currents were recorded across the full voltage range, quantification was standardised using the 20⍰mV pre-pulse step, which consistently produced robust and well-resolved tail currents across variants.

Recovery from inactivation (RFI) was evaluated using a paired-pulse protocol with inter-pulse intervals ranging from 10 to 1600⍰ms. The time required for 50% recovery (RRT₅₀) was extracted from normalized (P2/P1) current amplitudes.

Extracted parameters included peak current amplitude, current density (CD*), voltage dependence of activation/inactivation (V0.5), time-to-peak, Irel@50ms, τDeact, and RRT₅₀.

### Data Analysis

Data were analysed using MATLAB (The MathWorks, Inc.). Quality control thresholds and filtering criteria are described below: All recordings were subjected to rigorous quality control (QC) filtering to ensure data reliability and consistency across experiments. Inclusion criteria required a seal resistance >500 MΩ, cell capacitance (C_slow) between 3–50 pF, and series resistance <50 MΩ. Leak currents were corrected within a ±40 pA window. Functional benchmarks included a time-to-peak of <5 ms at 20 mV, peak activation currents between 200–5000 pA, and recovery-from-inactivation (RFI) amplitudes <400 pA.

Sample size calculations targeted 90% power to detect a 25% difference in functional metrics at α = 0.05. Data normality was assessed using the Kolmogorov–Smirnov test according to the sample sizes obtained. Non-Gaussian current density (CD*) values were square-root transformed and normalized to WT means per plate. Deactivation time constants (τ_Deact_) were log₁₀-transformed to improve distribution symmetry and enable parametric analysis ^22, 23^. We selected fixed transformations (square root and log₁₀) rather than adaptive methods requiring dataset-specific parameter estimation or arbitrary offsets for zero values. This approach is widely used in neurophysiology, and provides a simple, reproducible, and interpretable framework that ensures consistency across experiments while effectively reducing skewness. Outliers were removed using the ROUT algorithm (Q = 1%).

Functional classification was based on Z-scores relative to the distribution of 25 benign/likely benign (B/LB) control variants. Variants were considered functionally abnormal if their Z-scores exceeded ±2 standard deviations from the control mean; otherwise, they were classified as functionally normal. Concordance was defined as agreement between functional classification and ClinVar assertions (i.e., P/LP = abnormal; B/LB = normal). Sensitivity, specificity, and overall classification accuracy were calculated, and performance across metrics was compared using Fisher’s Exact Test.

To quantify the strength of functional evidence, Odds of Pathogenicity (OddsPath) values were calculated for each core metric in accordance with ClinGen Sequence Variant Interpretation (SVI) recommendations.

### Equations used in APC Analysis

Calculations were implemented in MATLAB during analysis.

*Current Density:*

Current Density (pA/pF) = current amplitude/capacitance

*Conductance:*

GNa = INa/(V – Vrev)

where GNa is the conductance, INa is the peak sodium current, V is the voltage and Vrev is the reversal potential. Conductance is normalized to the maximal conductance and fitted to the Boltzmann equation to calculate V_0.5_.

*Steady state inactivation:*

INa = INa / INaMax

where INa is the peak sodium current and INaMax is the maximal peak sodium conductance. Steady-state inactivation currents are normalized to the maximal current and fitted to the Boltzmann equation to calculate V_0.5_.

*Boltzmann equation:*

Boltzmann Equation = Imax + (Imax – Imin) / (1 + exp^((V0.5 - x)/k))

where Imax is the maximum peak current, Imin is the minimum peak current, V0.5 is the voltage of half-maximal activation or inactivation, and k is the slope factor.

*Recovery from inactivation:*

Recovery from inactivation current = P2/P1

where P1 is the peak at the control pulse and P2 is the peak at the test pulse occurring following increasing time intervals. This is fitted to a double exponential.

Double exponential = Y0+ ((Plateau-Y0)*(100-PercentFast)*.01)*(1-exp(-KSlow*x)) + ((Plateau-Y0)*PercentFast*.01)*(1-exp(-KFast*x))

### Multidimensional Data Visualization and Functional Severity Analysis

Variant phenotypes were quantified by comparing mean normalized current density (CD*, x-axis) and deactivation time constant (τ_Deact_⍰, y-axis). To provide an intuitive, scale=independent measure of overall functional deviation that integrates multiple phenotypic dimensions into a single metric, a Combined Z-score (CmbZ) was calculated as the Euclidean distance of individual parameter Z-scores:

CmbZ = SQRT [CD* Z^2^ + τ*_Deact_*’ Z^2^]

This composite metric captures cumulative deviation across orthogonal functional properties, channel output (CD*) and gating kinetics (τ_Deact_⍰), without assuming correlation, weighting, or shared variance between parameters. CmbZ is interpreted throughout as an effect⍰size metric reflecting functional severity, rather than as a test statistic or an indicator of statistical significance.

For variants in which only CD* could be reliably quantified, τ_Deact_⍰ was conservatively assigned a Z-score of zero for CmbZ calculation. This approach preserves interpretability by capturing unidimensional functional deviation while avoiding extrapolation beyond measured data.

### Bubble Plot Generation

Bivariate visualizations were generated using a custom Python script (v11.9) built on matplotlib and pandas. Variants were plotted in CD*–τ_Deact_⍰ space, with bubble area scaled to the Combined Z⍰score (CmbZ) and colour intensity reflecting the magnitude of composite deviation. Variants with low deviation (CmbZ <⍰2) were rendered as uniform blue markers, with higher values represented by progressively warmer colors up to magenta (CmbZ ≥ 2.0). These categories are descriptive and intended to aid visual interpretation.

### Handling of Missing Data

Variants lacking one of the two functional parameters were positioned in axis margins (“gutters”) rather than in the central bivariate field to avoid imputation. Variants with complete datasets were displayed as circular markers, while gutter⍰placed variants were shown as diamonds. Marginal points were slightly offset beyond the axis limits to maintain visual distinction without distorting spatial relationships.

### Integrated Functional Visualization (CmbZ)

CmbZ represents the Euclidean distance of CD* and τ_Deact_⍰ Z⍰scores and provides a scale⍰independent summary of multidimensional functional deviation. Because it does not rely on a statistical distributional assumption, CmbZ is used solely as an integrated visualization metric. In accordance with ACMG/ClinGen recommendations, PS3/BS3 evidence strength is derived exclusively from the calibrated single⍰parameter evidence (CD* and τ_Deact_⍰). CmbZ is not used for statistical inference or evidence assignment.

## Results

### Electrophysiological Profiling of CACNA1G Variants

To generate calibrated functional evidence for variant interpretation, we characterized Cav3.1-mediated currents from canonical wild-type (NM_018896.5) together with 25 curated benign/likely benign (B/LB) and 16 pathogenic/likely pathogenic (P/LP) *CACNA1G* variants associated with neurological phenotypes (Fig 1, Table S1). Functional parameters, including current density (CD), voltage dependence, and gating kinetics, were assessed using standard protocols as detailed in the Methods.

#### 1. Current Density

Current density, estimated from peak inward current at –20 mV (Fig 2A) and normalized to cell capacitance (pA/pF), reflects functional channel activity at the plasma membrane. Recordings from Cav3.1-WT (n = 1179 cells across 62 plates) revealed a non-Gaussian distribution of raw CD values (Fig 2B–C; KS distance = 0.09758, P < 0.0001, Kolmogorov–Smirnov (KS) test). Square root transformation (CD*) improved normality (Fig 2D–E; KS distance = 0.03438, P = 0.0023, Kolmogorov–Smirnov test), enabling standardized comparisons across variants.

A priori power analysis indicated that a minimum of 38 cells per variant was required to detect a 25% difference in CD* with 90% power with a 95% CI (Table S2). Using CD* values from the 25 B/LB variants, we established a reference distribution for biologically normal channel function and calculated Z-scores for all variants. Variants with Z-scores between −2 and +2 were considered within the normal range; scores between –2 and −4 or +2 and +4 indicated moderate deviation, whereas scores beyond ±4 were interpreted as severely dysfunctional. As shown in Figure 3A, 24 of 25 B/LB variants fell within the normal range, with B/LB-discordant Cav3.1-A1089T (c.3265G>A) marginally outside the set threshold (Z = −2.3). In contrast, 13 of 16 P/LP variants exhibited CD* Z-scores outside the normal range, consistent with impaired channel function. Notably, three P/LP-discordant variants (I161F, T867P, and S1799T) displayed CD* values within the normal range (Fig 3A, Table S3).

**Figure 3.**
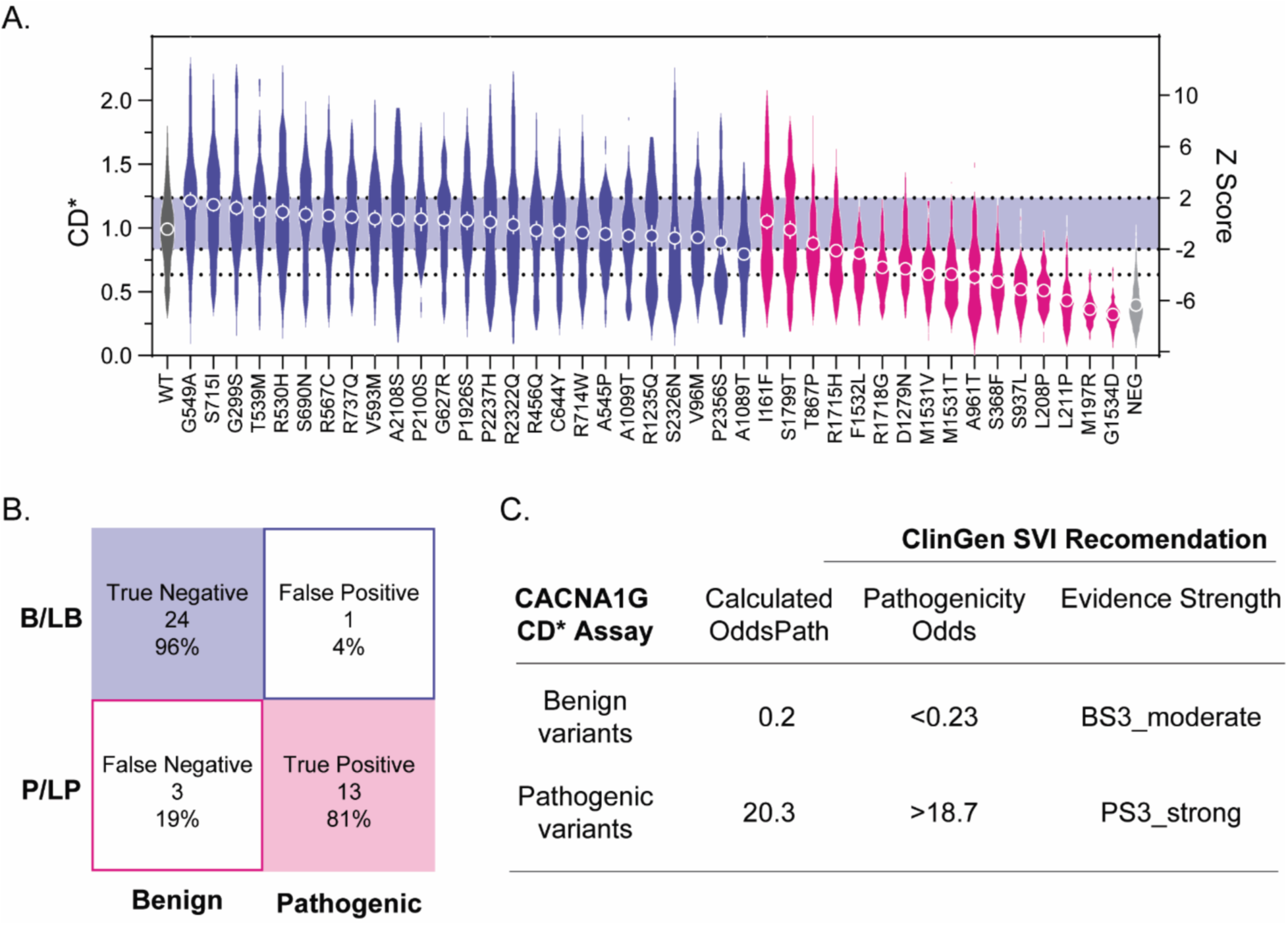
Current density analysis differentiates benign from pathogenic Cav3.1 variants with high accuracy. **A.** Square root–transformed current densities (CD*) for Cav3.1-WT and calibrating variants (left axis) with Z-scores (right y-axis) calibrated using 25 benign/likely benign variants. Normal function range is defined as the mean ± 2 standard deviations (SD) of benign CD*. B/LB (blue) cluster within Z-scores of −2 to +2 (shaded); whereas P/LP (magenta) predominantly fall outside this range. Horizontal shading denotes Z-scores within ±2; dotted lines indicate boundaries. Sample sizes for each variant are provided in Table S3. **B.** Confusion matrix summarizing CD*-based outcomes relative to reference variants. **C.** Under ClinGen Sequence Variant Interpretation guidelines, the calibrated CD* metric provides functional evidence supporting application of BS3_moderate and PS3_strong criteria.

To evaluate classification performance at the assay level, Odds of Pathogenicity (OddsPathPath) and Odds of Benignity (OddsPathBenign) were calculated from CD*-based classification outcomes. The CD* metric correctly classified 13 P/LP and 24 B/LB variants, yielding OddsPathPath = 20.3 and OddsPathBenign = 0.2 (Fig 3B, Table S4). In accordance with ClinGen Sequence Variant Interpretation guidelines, these values support use of CD* as primary functional evidence at the PS3_strong and BS3_moderate levels.^24^

#### 2. Cav3.1 Gating and Kinetics Properties

To determine whether additional functional abnormalities were present in variants with CD* values overlapping the normal range, we assessed gating and kinetic properties including voltage dependence of activation and inactivation (ActV₀.₅ and InacV₀.₅), activation kinetics (time to peak), extent of inactivation (Irel@50 ms), recovery from inactivation (RRT₅₀), and deactivation kinetics (τ_Deact_) were assessed to identify potential functional defects in variants with apparently normal current density (Fig 1). All measurements were obtained under standardized, automated conditions, to minimize experimenter bias and ensure that observed variability reflected intrinsic channel properties rather than methodological differences (see Methods).

Analysis of voltage dependent and kinetic parameters (Fig S1-S3) provided supportive functional context for variant interpretation. For example, P/LP variants with CD* values within the normal range, including T867P and S1799T, exhibited abnormal Z-scores for ActV₀.₅. Variant T867P additionally showed a hyperpolarizing shift in InacV₀.₅, whereas I161F remained within the B/LB reference range for both activation and inactivation parameters (Fig S1-S3). Recovery from inactivation was also significantly delayed for Cav3.1-S1799T compared with WT (Fig S2-S3). Complete numerical datasets for all gating and kinetic parameters are provided in Table S5 and S6.

Whereas most gating parameters showed limited discriminatory power when considered individually, deactivation kinetics (τ_Deact_) consistently differentiated pathogenic from benign reference variants and was therefore examined in greater detail.

##### 2.1 Deactivation kinetics

Deactivation time constants, τ_Deact_, were quantified by fitting a single exponential to tail current decay upon repolarization to –100 mV following 10 ms depolarizing steps from –120 mV to +50 mV (Fig 4). Quantification was performed following a pre-pulse to 20 mV, at which tail currents were consistently robust across variants. This analysis was feasible for 13 of the 16 P/LP variants; the remaining three variants (M197R, L211P, and G1534D) generated insufficient current for reliable fitting (Table S3).

**Figure 4.**
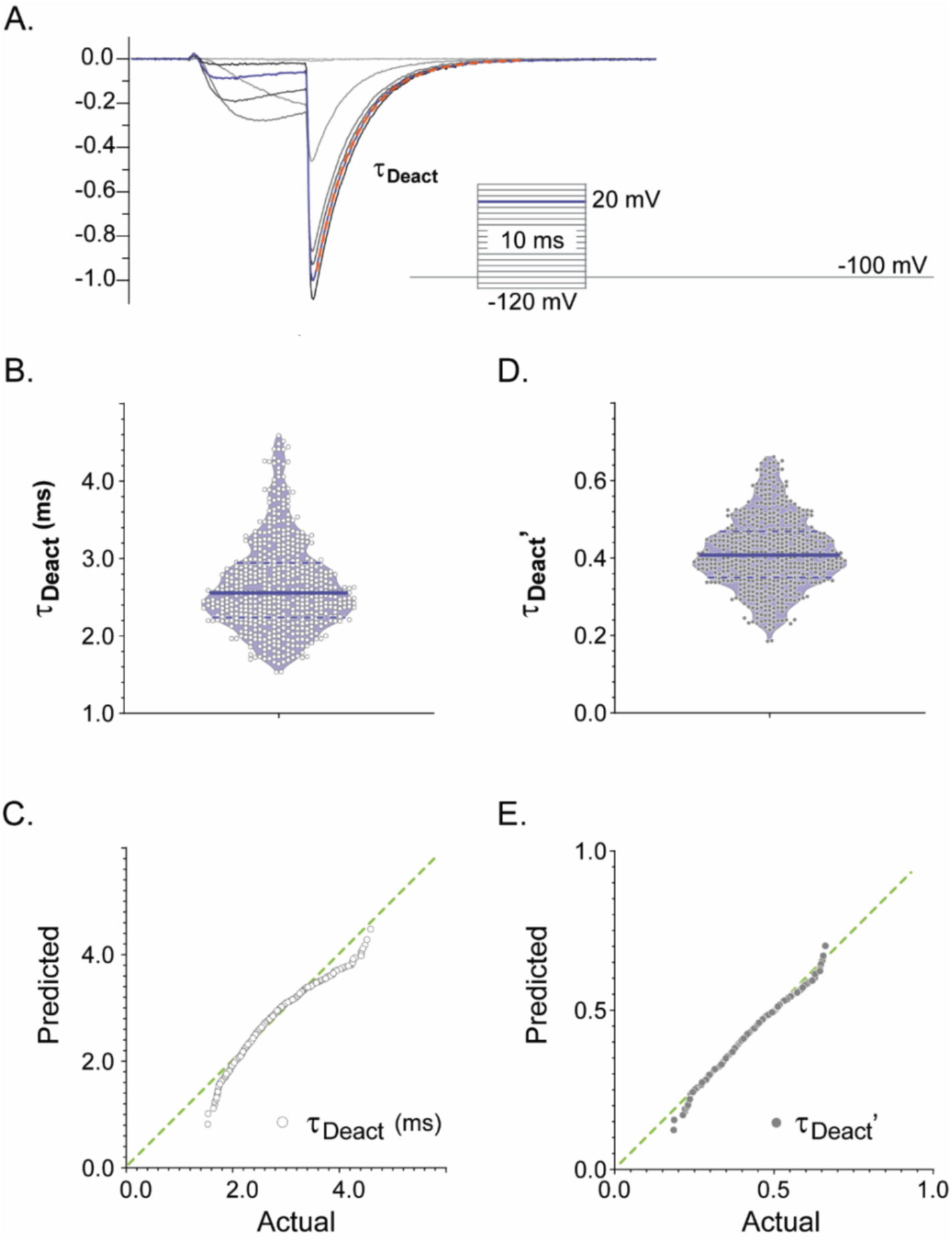
Distribution of Cav3.1-WT deactivation time constants (t_Deact_). **A** Representative tail current traces recorded upon repolarization to −100⍰mV following 10⍰ms depolarizing pre-pulses from −120⍰mV to +20⍰mV in 10⍰mV increments (inset). Deactivation kinetics were quantified from the tail current decay following a 20⍰mV pre-pulse (highlighted in blue), by fitting a single exponential function (orange) to extract t_Deact_. **B–C.** Violin and quantile–quantile (Q-Q) plots of raw Cav3.1-WT t_Deact_ values (n = 558 cells, 43 experiments), showing significant deviation from normality (KS distance = 0.08210, p < 0.0001). **D–E.** Violin and Q-Q plots of log_10_-transformed t_Deact_ (t_Deact_‘), demonstrating conformity to a Gaussian distribution (KS distance = 0.04000, p = 0.0332). Median values (solid blue line) and interquartile ranges (dotted blue lines) are indicated.

Because Z-score analysis assumes normally distributed data, raw τ_Deact_ values were log-transformed (τ_Deact_‘, see Methods) to improve normality and stabilize variance prior to statistical analysis. Log transformation substantially reduced skew and improved distribution symmetry (raw τ_Deact_ KS distance = 0.0821, p < 0.0001 vs τ_Deact_‘ KS distance = 0.04, p = 0.0332), enabling more reliable application of Z-scoring (Fig 4). *A priori* power analysis indicated that a minimum of 17 cells per variant was required to detect a 25% difference in τ_Deact_‘ with 90% power with a 95% CI (Table S2).

τ_Deact_‘ separated benign and pathogenic reference distributions (Fig 5A, Table S7). Z-score classification based on deactivation kinetics correctly classified 11 P/LP and 24 B/LB variants, yielding OddsPathPath = 21.2 and OddsPathBenign = 0.16 (Fig 5B, Table S4). Notably, P/LP variants S368F, T867P, M1531T, and S1799T exhibited marked alterations in deactivation kinetics, with S368F displaying faster deactivation and the remaining variants showing slower deactivation (Fig 5A, S3). These results indicate that τ_Deact_ capture gating-specific dysfunction not evident from current density analysis alone, providing complementary functional evidence for variant interpretation.

**Figure 5.**
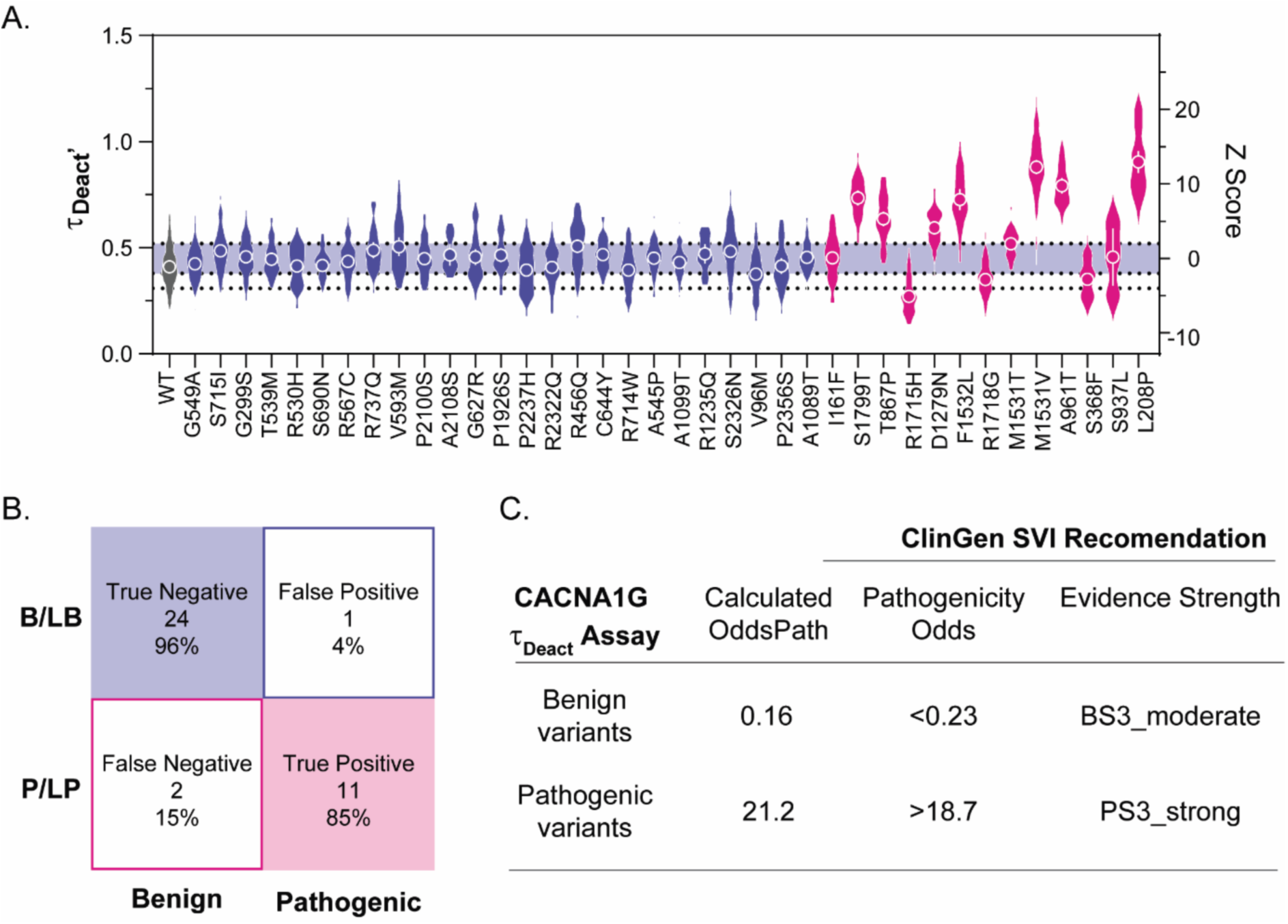
Deactivation kinetics provide supportive functional evidence for Cav3.1 variant interpretation. **A.** Deactivation time constants (τ_Deact_’) for WT Cav3.1 and variant channels (left y-axis), with corresponding Z-scores (right y-axis) calibrated using 25 benign/likely benign (B/LB) variants. Normal functional range is defined as mean ± 2 SD of B/LB τ_Deact_ values. B/LB variants (blue) cluster within Z-scores of −2 to +2 (shaded region), whereas P/LP variants (magenta) predominantly fall outside this range. Horizontal shading denotes Z-scores within ±2 and dotted lines indicate outliers. Variant-specific sample sizes in Table S7. **B.** Confusion matrix summarizing τ_Deact_‘-based outcomes relative to reference variants. **C.** Consistent with ClinGen Sequence Variant Interpretation guidelines, τ_Deact_‘ contributes supportive functional evidence for variant interpretation, whereas CD* serves as the primary metric supporting BS3_moderate and PS3_strong criteria.

### Application of the Calibrated Framework to VUS

To evaluate the clinical interpretability of calibrated functional evidence, we applied primary functional metrics, current density and deactivation kinetics, to variants of uncertain significance. In accordance with ClinGen SVI recommendations, each functional evidence was evaluated independently, and only the strongest applicable functional evidence was considered to avoid double counting. This framework was applied to five *CACNA1G* VUS identified in Australian patients with neurological phenotypes (Table S8).

Three variants (S1098N, A2126S, and R2149Q) exhibited functional profiles within the benign/likely benign (B/LB) reference range for both CD* and τ_Deact_’ (Fig 6 and Table S3, S7), indicating Cav3.1 channel function consistent with the benign reference distribution. In contrast, two VUS, R186Q and R1394Q, fell outside normal thresholds and displayed distinct functional abnormalities (Fig 6A, B). Variant R186Q showed a moderate reduction in CD* (Z = – 3.6) together with markedly prolonged τ_Deact_’ (Z = 6.3) (Fig, 6C). This variant also exhibited left-shifted voltage dependence of activation and inactivation (Fig 7A-B), delayed recovery from inactivation (Fig 7D), and otherwise normal activation and inactivation kinetics (Fig 7C and Fig S4). Variant R1394Q displayed a CD* values overlapping the B/LB range but markedly accelerated deactivation (Z = –10.2, Fig 6A, C). This phenotype was accompanied by right-shifted voltage dependence (Fig 7A, B), faster inactivation (Z= –3.0, Fig 7C), and within normal recovery and activation kinetics (Fig7D and Fig S4, respectively). Based on integration of calibrated functional evidence with existing clinical and genetic data, both R186Q and R1394Q met criteria supporting reclassification as likely pathogenic under ACMG/AMP guidelines (PS3; Table S1).

**Figure 6.**
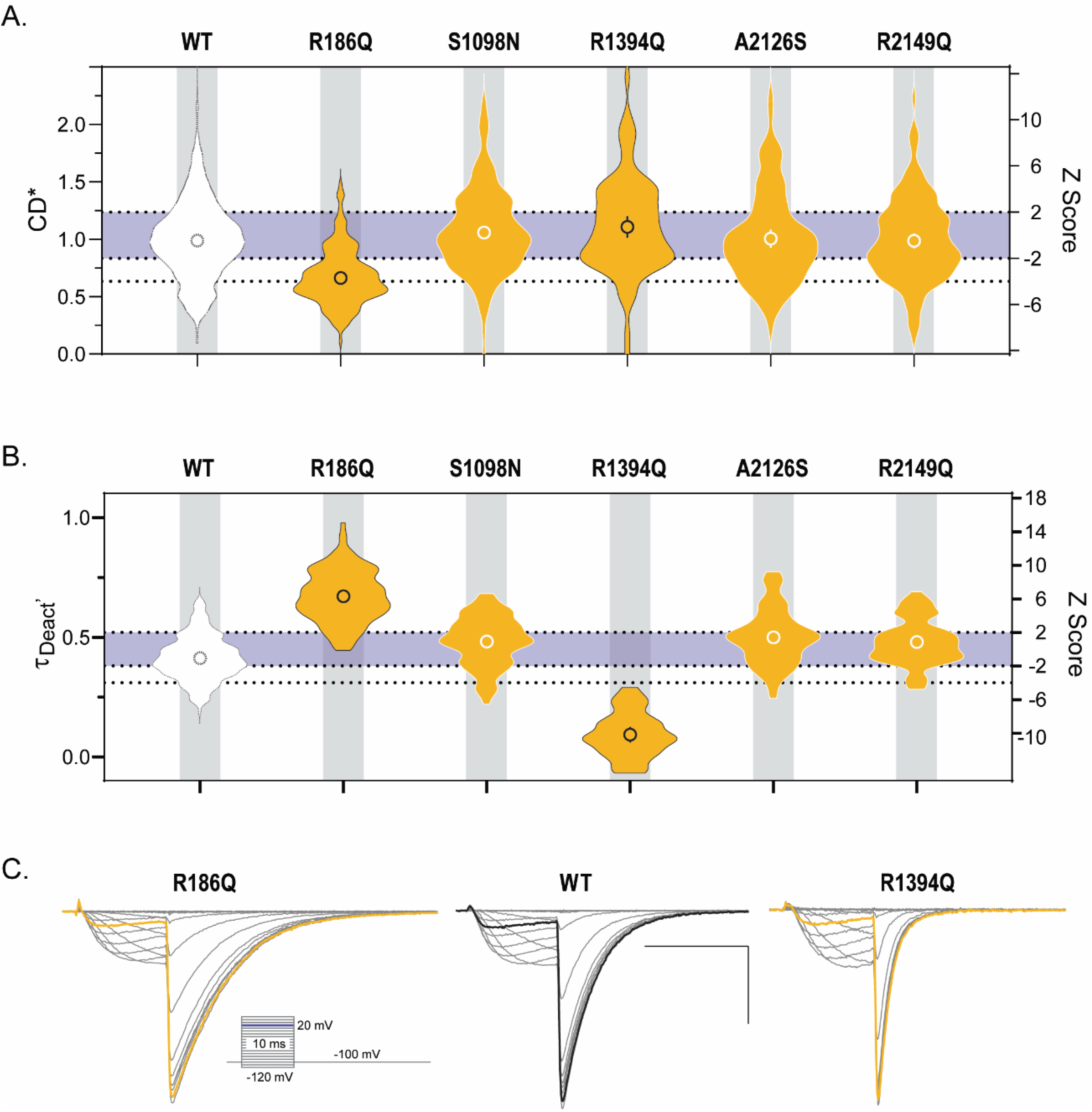
Integrated CD* and τ_Deact_’ assessment supports interpretation of CACNA1G VUS. **A.** Mean ± 95% confidence intervals (CI) for CD* of WT and VUS channels (left y-axis), with corresponding Z-scores (right y-axis). **B.** Mean ± 95% CI for τ_Deact_’ of WT and VUS channels (left y-axis), with corresponding Z-scores (right y-axis). In panels **A** and **B**, the normal functional range defined as mean ± 2 SD of B/LB variants, is shaded in blue. **C.** Representative tail current traces following a 10 ms activating pre-pulse (inset) for WT and selected VUS, illustrating domain-specific gating alterations. DI-S4 variant R186Q shows slower deactivation, whereas DIII-S4 variant R1394Q exhibits faster deactivation. Traces used for quantification are highlighted in dark grey (WT) and yellow (VUS). Calibration bars: 10 ms and 200 pA. Variant-specific sample sizes are provided in Tables S3 and S7.

**Figure 7.**
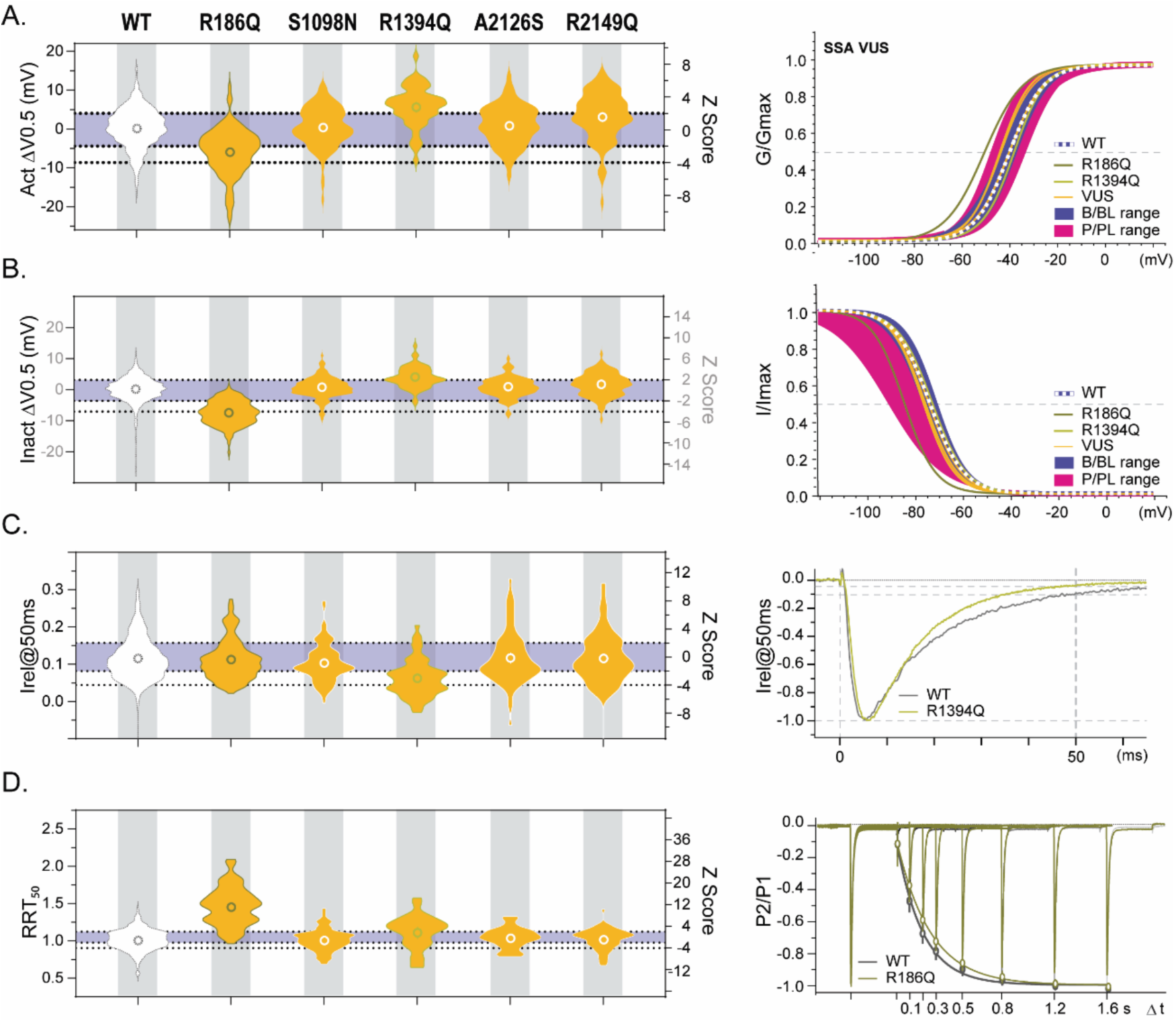
Voltage dependence and kinetic parameters provide supportive functional context for interpretation of CACNA1G VUS. **A.** Left: Relative voltage dependence of activation (act ΔV₀.₅). Right: Normalized conductance–voltage plots showing the normal range (B/LB, magenta) and abnormal range (P/LP, blue) for activation. **B.** Left: Relative voltage dependence of half-inactivation (inact ΔV₀.₅). Right: Current availability plots showing the normal (B/LB, magenta) and abnormal (P/LP, blue) ranges for inactivation. In panels **A–B**, WT is shown as dashed white lines, VUS within the B/LB range are shown in yellow/white, and VUS outside this range in yellow/black. **C.** Left: Inactivation (Irel@50ms). Right: Representative traces for WT (dark gray) and R1394Q (yellow). **D.** Left: Recovery from inactivation (Relative Recovery Time, RRT₅₀), defined as the ratio of time for 50% recovery in a variant relative to WT. Right: Representative traces for WT (dark gray) and R186Q (yellow). For panels **A–D**, mean ± 95% CI are shown on the left y-axis and Z-scores on the right y-axis. Blue shading denotes Z-scores within ±2, and dotted lines indicate values beyond this range. Variant-specific sample sizes are provided in Tables S5 and S6.

### Multidimensional visualization functional severity across CACNA1G variants

Joint visualization of CD* and τ_Deact_’ provides an integrated view of *CACNA1G* variant effects that complements single-metric analyses. Figure 8 implements this bivariate representation by projecting channel output (CD*) and gating kinetics (τ_Deact_’) into a shared feature space. Variant positions are mapped onto the Cav3.1 protein schematic (Fig 8A), and CD* values are plotted against τ_Deact_’ in Fig 8B. Bubble color and size reflect the Combined Z-score (CmbZ; see Methods), which summarizes multidimensional deviation and is used exclusively for visualization.

**Figure 8.**
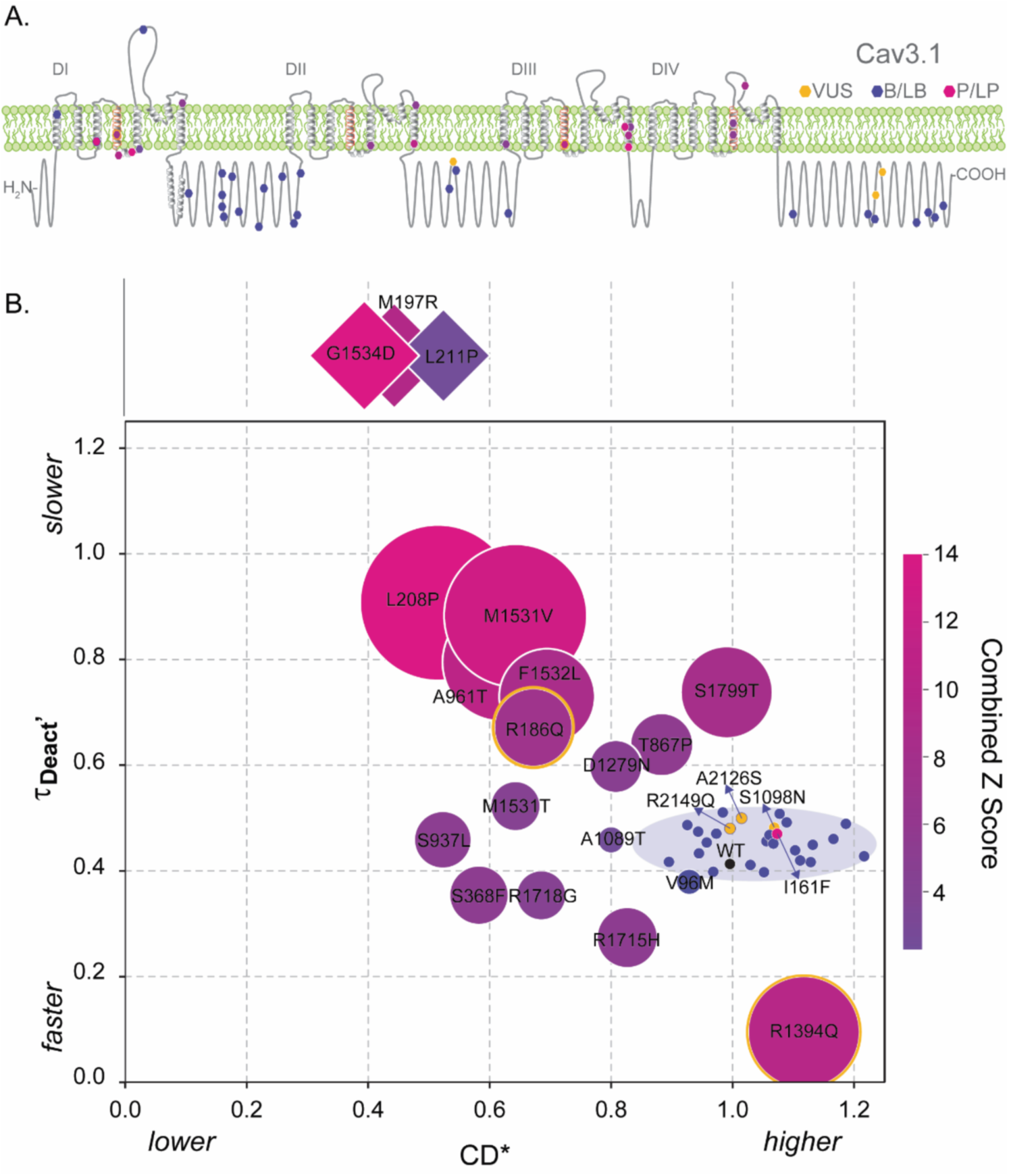
Bivariate visualization of CACNA1G variant effects on current density and deactivation kinetics. **A.** Schematic Cav3.1 schematic indicating the positions of variants with B/LB and P/LP functional profiles (colored as in **B.**). **B.** Bubble plot of mean current density (CD*, x-axis) vs. deactivation time constant (τ_Deact_’, y-axis) for all reported variants. Circles mark variants with both parameters quantified. Colour and size represent the Combined Z-score (CmbZ, see Methods). Composite deviation is mapped from blue (low; CmbZ < 2.0) to magenta (high; CmbZ ≥ 2.0). The shaded ellipse marks distances from the B/LB centroid in Z⍰space; points inside fall within −2 ≤ CmbZ ≤ 2. Bubble colour/size illustrate CmbZ purely as an integrated effect⍰size visualization; CmbZ is not used to assign PS3/BS3 strength. High-impact P/LP variants lacking measurable τ_Deact_’ (Cav3.1-M197R, -L211P, and -G1534D) appear as diamond markers positioned along the CD* axis and offset into the axis gutters to retain representation. B/LB variants are shown in blue; Cav3.1-I161F, a P/LP variant with normal function, appears in magenta. VUS are highlighted in yellow, with marker outline or fill indicating P/LP-like or B/LB-like function, respectively.

Across the calibrated dataset, variants span a continuum from WT⍰like behavior to substantial functional perturbation. B/LB reference variants cluster tightly near the Cav3.1-WT centroid, whereas most P/LP variants deviate markedly through reduced channel output, altered deactivation kinetics, or both. High⍰impact P/LP variants lacking measurable τ_Deact_⍰ are displayed along the CD* axis. Cav3.1⍰I161F, despite its P/LP annotation, resides within the WT⍰like cluster, illustrating how calibrated functional evidence can resolve discordant classification. Several VUS map near the benign reference cluster, whereas R186Q and R1394Q show clear multidimensional deviation consistent with pathogenic-like dysfunction.

This multidimensional representation contextualizes graded functional effects across the *CACNA1G* variant landscape. In accordance with ACMG/ClinGen guidance, assignment of PS3 or BS3 evidence strength is derived exclusively from calibrated single-parameter metrics (CD* and τ_Deact⍰), whereas CmbZ serves solely as an integrative, descriptive visualization and is not used for statistical inference or evidence assignment.

## Discussion

This study establishes a calibrated functional framework for interpretation of *CACNA1G* variants of uncertain significance (VUS), based on systematic functional profiling of 25 known benign/likely benign and 16 known pathogenic/likely pathogenic variants. Using automated patch-clamp electrophysiology, we defined reference distributions for multiple biophysical properties of Cav3.1 channel function. Among the parameters assessed, normalized current density and deactivation kinetics provided the most consistent and discriminatory functional signals. When evaluated independently and in accordance with ACMG/ClinGen recommendations, these calibrated metrics generated functional evidence supporting application of PS3 (strong) and BS3 (moderate) criteria. Importantly, this framework enabled resolution of VUS with functional profiles overlapping either the benign or pathogenic reference distributions, providing evidence to support variant reclassification within established variant interpretation guidelines. Collectively, these findings demonstrate the value of combining complementary electrophysiological metrics to support conservative clinical interpretation of CACNA1G variants Normalized current density, reflecting the functional availability of Cav3.1 channels at the plasma membrane, showed strong discriminatory power between B/LB and P/LP variants. More than 80% of P/LP variants exhibited CD* values below the benign range, consistent with reduced channel abundance or stability. Deactivation kinetics provided complementary insight by revealing gating abnormalities in two of the three pathogenic variants whose CD* values overlapped the benign range. These observations indicate that assessment of channel gating, in addition to current amplitude, can identify functionally relevant defects not captured by single-metric analysis.

Deactivation kinetics represent a physiologically relevant and technically robust measure of Cav3 channel function, reflecting intrinsic channel closure during action potential repolarization. Unlike activation or inactivation protocols, which often require non-physiological voltage steps or prolonged conditioning pulses and can be sensitive to artifacts such as series-resistance errors or current rundown, deactivation can be quantified using brief depolarization steps that more closely approximate physiological conditions. For T-type calcium channels, appropriate deactivation ensures transient Ca²⁺ influx that supports rebound depolarization and rhythmic burst firing without promoting excessive afterdepolarization.^25^ Disruption of this balance may alter neuronal oscillatory behaviour and has been implicated in pathological rhythms, including those associated with absence epilepsy.^26–28^

Among P/LP control variants examined, the previously reported gain-of-function variants A961T^9^ and M1531V^8^ clustered together in our bivariate analysis. M197R also exhibited a severe loss-of-function phenotype. In contrast, Cav3.1-I161F, located in domain I near the S2–S3 linker and the cytoplasmic end of S3 (Fig 1A, 8A), has conflicting assertions in ClinVar (Variation ID: 1027470; Likely pathogenic; Uncertain significance). Despite its position within a structurally sensitive region, I161F displayed electrophysiological properties indistinguishable from wild type, albeit with greater variability in some parameters. Although this platform lacks neuronal contextual cues, the data indicate that I161F does not meaningfully alter Cav3.1 channel function under the conditions tested. These observations underscore the importance of experimental functional validation in resolving conflicting variant classifications and highlight the value of calibrated electrophysiology in refining variant interpretation.

Mechanistically, both VUS with abnormal functional profiles, R186Q and R1394Q, result in neutralization of arginine residues within the S4 segment of voltage-sensing domains I and III, respectively (Fig 8A). Because S4 positive charges are essential for detecting changes in membrane potential and initiating conformational transitions that drive channel gating, their loss is known to impair voltage sensitivity and gating kinetics in voltage-gated channels.^29–31^ The observed shifts in activation and inactivation, altered deactivation kinetics in both variants, and reduced current density in R186Q are consistent with disrupted voltage-sensor function. These findings underscore the relevance of CD* and τ_Deact_’ as sensitive readouts of voltage-sensor integrity and support their use as core metrics for variant classification.

The dual-metric framework described here enhances sensitivity while maintaining specificity when incorporating functional data into *CACNA1G* variant interpretation. In alignment with ClinGen Sequence Variant Interpretation guidance, CD* and τ_Deact_’ were evaluated independently, and and only the strongest applicable functional evidence was considered to avoid double counting. Additional parameters, including voltage dependence, activation and inactivation kinetics, and recovery from inactivation, provided mechanistic context in selected cases but exhibited greater overlap between benign and pathogenic reference distributions and therefore limited discriminatory utility at the population level. Accordingly, these parameters were not used for formal evidence strength assignment. This calibrated, evidence restricted approach supports reproducible integration of functional data into clinical variant interpretation while maintaining conservative interpretive standards.

When generated in specialized functional laboratories, calibrated electrophysiological measurements such as those described here can contribute external functional evidence for consideration during CACNA1G variant interpretation, particularly for missense variants that remain unresolved following standard clinical, genetic, and computational assessment.

Despite its scalability and standardization, high-throughput automated patch-clamp performed in HEK293 cells have intrinsic limitations. HEK293 cells are kidney-derived cells lacking the neuronal molecular environment, accessory proteins, and activity-dependent regulation that influence Cav3.1 channel behavior *in vivo*. As a result, electrophysiological measurements obtained in this system may not fully capture context-dependent modulation of channel function present in native neuronal circuits.

In addition, biophysical characterization in a heterologous expression system does not directly predict clinical phenotype. Gain- or loss-of-function effects may be buffered, amplified, or modified by neuronal network properties, developmental stage, or an individual’s broader genetic background. Accordingly, the functional effects quantified here should be interpreted as component evidence rather than as stand-alone predictors of disease. These limitations underscore the necessity of integrating calibrated functional data with clinical, genetic, and computational evidence within established variant-interpretation frameworks.

In summary, we describe a calibrated functional framework for evaluating *CACNA1G* missense variants, with normalized current density and deactivation kinetics identified as core discriminatory metrics. Application of this framework to five variants of uncertain significance revealed functional abnormalities for R186Q and R1394Q, consistent with impaired Cav3.1 channel behavior, whereas S1098N, A2126S, and R2149Q exhibited functional profiles overlapping the benign reference range.

In the clinical context, functional evidence generated using this framework was incorporated alongside genetic and clinical information during variant interpretation, contributing to reassessment of R186Q and R1394Q as likely pathogenic. By contrast, variants with WT-like functional profiles aligned with clinical features not suggestive of *CACNA1G*-driven disease. These findings illustrate how calibrated electrophysiological data can inform interpretation of unresolved *CACNA1G* variants when applied conservatively within ACMG/ClinGen guidelines.

Overall, this work provides a reproducible approach for generating standardized functional evidence applicable to clinical variant interpretation. By supporting resolution of selected *CACNA1G* missense variants that remain uncertain after standard evaluation, this framework promotes more informed and cautious use of functional data in precision medicine for Cav3.1-associated neurological disorders.

## Acknowledgements

We thank Bradley Wakefield and Marijka Batterham of the Statistical Consulting Centre at the University of Wollongong for their statistical guidance and support. We also acknowledge Jeffrey R. McArthur (PhD) for his assistance with Python coding. This work was supported by the Medical Research Future Fund, Genomics Health Futures Mission grants: MRF2016760 (DJA/JIV/CAN) and MRF2007677 (PJL).

## Author Contributions

Conceptualization: RKF, JIV, CAN; data curation: RKF, CYT, NM, PJL, PS, AM, BAT, GG, HG, CAN; formal analysis: RKF, CYT, NM, JGM, JIV, CAN; funding acquisition: PJL, DJA, JIV, CAN; methodology: RKF, DJA, JIV, CAN; project administration: RKF, JIV, CAN; resources: LRG, DJA, JIV; supervision: LRG, DJA, JIV, CAN; writing—original draft: RKF, JIV, CAN; writing—review and editing: RKF, CYT, NM, JGM, PJL, PS, AM, BAT, GG, HG, LRG, DJA, JIV, CAN. All authors read and approved the final manuscript.

## Data Availability

All data generated or analyzed during this study are included in this published article and its supplementary information files. The datasets used and/or analyzed during the current study are available from the corresponding author on reasonable request.

## References

1. Landrum MJ, Chitipiralla S, Kaur K, et al. ClinVar: updates to support classifications of both germline and somatic variants. Nucleic Acids Res. 2025 Jan 6;53(D1):D1313–D21.

2. Richards S, Aziz N, Bale S, et al. Standards and guidelines for the interpretation of sequence variants: a joint consensus recommendation of the American College of Medical Genetics and Genomics and the Association for Molecular Pathology. Genet Med. 2015 May;17(5):405–24.

3. Son H, Yoon JG, Kim MJ, Moon J, Kim HJ. First Cases of Spinocerebellar Ataxia 42 in Two Korean Families. J Mov Disord. 2023 Jan;16(1):110–3.

4. Hashiguchi S, Doi H, Kunii M, et al. Ataxic phenotype with altered Ca(V)3.1 channel property in a mouse model for spinocerebellar ataxia 42. Neurobiol Dis. 2019 Oct;130:104516.

5. Morino H, Matsuda Y, Muguruma K, et al. A mutation in the low voltage-gated calcium channel CACNA1G alters the physiological properties of the channel, causing spinocerebellar ataxia. Mol Brain. 2015 Dec 29;8:89.

6. Calhoun JD, Hawkins NA, Zachwieja NJ, Kearney JA. Cacna1g is a genetic modifier of epilepsy in a mouse model of Dravet syndrome. Epilepsia. 2017 Aug;58(8):e111–e5.

7. Calhoun JD, Hawkins NA, Zachwieja NJ, Kearney JA. Cacna1g is a genetic modifier of epilepsy caused by mutation of voltage-gated sodium channel Scn2a. Epilepsia. 2016 Jun;57(6):e103–7.

8. Davakan A, Cmarko L, Ribeiro Oliveira-Mendes B, et al. Electrophysiological classification of CACNA1G gene variants associated with neurodevelopmental and neurological disorders. Front Pharmacol. 2025;16:1613072.

9. Qebibo L, Davakan A, Nesson-Dauphin M, et al. The characterization of new de novo CACNA1G variants affecting the intracellular gate of Cav3.1 channel broadens the spectrum of neurodevelopmental phenotypes in SCA42ND. Genet Med. 2025 Mar;27(3):101337.

10. Weiss N, Zamponi GW. Genetic T-type calcium channelopathies. J Med Genet. 2020 Jan;57(1):1–10.

11. Anderson MP, Mochizuki T, Xie J, et al. Thalamic Cav3.1 T-type Ca2+ channel plays a crucial role in stabilizing sleep. Proc Natl Acad Sci U S A. 2005 Feb 1;102(5):1743–8.

12. Yabuki Y, Hori K, Zhang Z, et al. Cav3.1 T-Type Calcium Channel Acts as a Gateway for GABAergic Excitation in the Medial Prefrontal Cortex That Leads to Chronic Psychological Stress Responses in Mice. Acta Physiol (Oxf). 2025 May;241(5):e70043.

13. Berecki G, Helbig KL, Ware TL, et al. Novel Missense CACNA1G Mutations Associated with Infantile-Onset Developmental and Epileptic Encephalopathy. Int J Mol Sci. 2020 Aug 31;21(17).

14. Ma JG, O’Neill MJ, Richardson E, et al. Multisite Validation of a Functional Assay to Adjudicate SCN5A Brugada Syndrome-Associated Variants. Circ Genom Precis Med. 2024 Aug;17(4):e004569.

15. Thomson KL, Jiang C, Richardson E, et al. Clinical interpretation of KCNH2 variants using a robust PS3/BS3 functional patch-clamp assay. HGG Adv. 2024 Apr 11;5(2):100270.

16. Ng CA, O’Neill MJ, Padigepati SR, et al. Calibrated Functional Data Decrease Clinical Uncertainty for KCNH2-Related Long-QT Syndrome. Circ Genom Precis Med. 2025 Aug;18(4):e005204.

17. Vanoye CG, Desai RR, Fabre KL, et al. High-Throughput Functional Evaluation of KCNQ1 Decrypts Variants of Unknown Significance. Circ Genom Precis Med. 2018 Nov;11(11):e002345.

18. O’Neill MJ, Ma JG, Aldridge JL, et al. Automated patch clamp data improve variant classification and penetrance stratification for SCN5A-Brugada syndrome. Eur Heart J. 2025 Nov 18.

19. Glazer AM, Wada Y, Li B, et al. High-Throughput Reclassification of SCN5A Variants. Am J Hum Genet. 2020 Jul 2;107(1):111–23.

20. Ruano L, Melo C, Silva MC, Coutinho P. The global epidemiology of hereditary ataxia and spastic paraplegia: a systematic review of prevalence studies. Neuroepidemiology. 2014;42(3):174–83.

21. Ng CA, Farr J, Young P, et al. Heterozygous KCNH2 variant phenotyping using Flp-In HEK293 and high-throughput automated patch clamp electrophysiology. Biol Methods Protoc. 2021;6(1):bpab003.

22. West RM. Best practice in statistics: The use of log transformation. Ann Clin Biochem. 2022 May;59(3):162–5.

23. Curran-Everett D. Explorations in statistics: the log transformation. Adv Physiol Educ. 2018 Jun 1;42(2):343–7.

24. Brnich SE, Abou Tayoun AN, Couch FJ, et al. Recommendations for application of the functional evidence PS3/BS3 criterion using the ACMG/AMP sequence variant interpretation framework. Genome Med. 2020 Dec 31;12(1):3.

25. Zamponi GW, Lory P, Perez-Reyes E. Role of voltage-gated calcium channels in epilepsy. Pflugers Arch. 2010 Jul;460(2):395–403.

26. Lee SE, Lee J, Latchoumane C, et al. Rebound burst firing in the reticular thalamus is not essential for pharmacological absence seizures in mice. Proc Natl Acad Sci U S A. 2014 Aug 12;111(32):11828–33.

27. Maksemous N, Blayney CD, Sutherland HG, et al. Investigation of CACNA1I Cav3.3 Dysfunction in Hemiplegic Migraine. Front Mol Neurosci. 2022;15:892820.

28. Gobbo D, Scheller A, Kirchhoff F. From Physiology to Pathology of Cortico-Thalamo-Cortical Oscillations: Astroglia as a Target for Further Research. Front Neurol. 2021;12:661408.

29. Bezanilla F. The voltage sensor in voltage-dependent ion channels. Physiol Rev. 2000 Apr;80(2):555–92.

30. Swartz KJ. Sensing voltage across lipid membranes. Nature. 2008 Dec 18;456(7224):891–7.

31. Catterall WA. Ion channel voltage sensors: structure, function, and pathophysiology. Neuron. 2010 Sep 23;67(6):915–28.

