## Supplemental Table 1 for "Calibrated high-throughput electrophysiology enables clinical interpretation of *CACNA1G* missense variants"

**Table S1: *CACNA1G* Variant controls and VUS**

**Benign controls**

| Gene | Transcript | Genomic Location and Change | DNA Change | Protein Change | gnomAD AF | | | gnomAD v4.1.0 Grpmax Filtering AF | gnomAD homozygotes | Clinvar Classification (19 Jan 26) | ACMG/AMP Criteria | Final Classification |
|---|---|---|---|---|---|---|---|---|---|---|---|---|
| | | | | | V2.1.1 (Exomes) | v3.1.2 | v4.1.0 (Total) | | | | | |
| CACNA1G | NM_018896.5 | chr17(GRCh38):g.50568913G>A | c.286G>A | p.Val96Met | 3.82E-04 | 3.02E-04 | 6.59E-04 | 0.0007717 European (non-Finnish) | 0(v2.1.1), 0(v3.1.2), 0(v4.1.0) | 1234274 Benign/Likely benign | PP2, PP3 (M), BS1, BS2 (P) | Likely Benign |
| CACNA1G | NM_018896.5 | chr17(GRCh38):g.50572702G>A | c.895G>A | p.Gly299Ser | 1.87E-03 | 1.52E-03 | 1.86E-03 | 0.001972 European (non-Finnish) | 1(v2.1.1),1(v3.1.2), 7(v4.1.0) | 720406 Benign/Likely benign | PP2, BS1, BS2 (S), BP4 | Benign |
| CACNA1G | NM_018896.5 | chr17(GRCh38):g.50575769G>A | c.1367G>A | p.Arg456Gln | 6.88E-03 | 7.03E-03 | 9.91E-03 | 0.01184 European (non-Finnish) | 8(v2.1.1),12(v3.1.2), 108(v4.1.0) | 771920 Benign | BS1, BS2 (S), PP1 (P) | Benign |
| CACNA1G | NM_018896.5 | chr17(GRCh38):g.50575991G>A | c.1589G>A | p.Arg530His | 1.44E-03 | 4.15E-03 | 8.48E-04 | 0.01465 African/African American | 5(v2.1.1), 5(v3.1.2), 14(v4.1.0) | 790179 Benign/Likely benign | BS1, BS2 (S), BP4 | Benign |
| CACNA1G | NM_018896.5 | chr17(GRCh38):g.50576018C>T | c.1616C>T | p.Thr539Met | 4.12E-04 | 1.77E-04 | 7.04E-05 | 0.001345 East Asian | 0(v2.1.1), 0(v3.1.2), 0(v4.1.0) | 2176332 Benign | BS1, BS2 (P), BP4 | Likely Benign |
| CACNA1G | NM_018896.5 | chr17(GRCh38):g.50576035G>C | c.1633G>C | p.Ala545Pro | 4.46E-04 | 4.40E-04 | 2.35E-04 | 0.00006066 Admixed American | 0(v2.1.1), 0(v3.1.2), 2(v4.1.0) | 2152746 Benign/Likely benign | BS1, BS2 (P), BP4 | Likely Benign |
| CACNA1G | NM_018896.5 | chr17(GRCh38):g.50576048G>C | c.1646G>C | p.Gly549Ala | 5.01E-04 | 6.83E-04 | 3.21E-04 | 0.003444 Middle Eastern | 0(v2.1.1), 0(v3.1.2), 1(v4.1.0) | 733508 Benign/Likely benign | BS1, BS2 (P), BP4 | Likely Benign |
| CACNA1G | NM_018896.5 | chr17(GRCh38):g.50576101C>T | c.1699C>T | p.Arg567Cys | 3.01E-03 | 9.39E-03 | 1.76E-03 | 0.03345 African/African American | 8(v2.1.1), 19(v3.1.2), 36(v4.1.0) | 777202 Benign/Likely benign | BS1, BS2 (S), PP3 (P) | Benign |
| CACNA1G | NM_018896.5 | chr17(GRCh38):g.50576179G>A | c.1777G>A | p.Val593Met | 3.31E-04 | 2.50E-04 | 1.93E-04 | 0.0004039 Admixed American | 1(v2.1.1), 0(v3.1.2), 1(v4.1.0) | 2057114 Benign/Likely benign | BS1, BS2 (S), BP4 | Benign |
| CACNA1G | NM_018896.5 | chr17(GRCh38):g.50576281G>A | c.1879G>A | p.Gly627Arg | 1.36E-03 | 7.62E-04 | 1.02E-03 | 0.004906 South Asian | 1(v2.1.1), 0(v3.1.2), 3(v4.1.0) | 718649 Benign/Likely benign | BS1, BS2 (S), BP4 | Benign |
| CACNA1G | NM_018896.5 | chr17(GRCh38):g.50578194G>A | c.1931G>A | p.Cys644Tyr | 5.70E-04 | 4.34E-04 | 4.08E-04 | 0.001878 Middle Eastern | 0(v2.1.1), 0(v3.1.2), 0(v4.1.0) | 634620 Benign/Lilely benign/VUS | BS1, BS2 (P), BP4 | Likely Benign |
| CACNA1G | NM_018896.5 | chr17(GRCh38):g.50578332G>A | c.2069G>A | p.Ser690Asn | 2.60E-04 | 8.02E-04 | 1.47E-04 | 0.002426 African/African American | 0(v2.1.1), 0(v3.1.2), 0(v4.1.0) | 719545 Benign/Likely benign | BS1, BS2 (P) BP4 | Likely Benign |
| CACNA1G | NM_018896.5 | chr17(GRCh38):g.50578403C>T | c.2140C>T | p.Arg714Trp | 2.36E-04 | 3.81E-04 | 1.21E-04 | 0.0007705 African/African American | 0(v2.1.1), 0(v3.1.2), 0(v4.1.0) | 2203297 Likely benign | BS1, BS2 (P), BP4 | Likely Benign |
| CACNA1G | NM_018896.5 | chr17(GRCh38):g.50578407G>T | c.2144G>T | p.Ser715Ile | 2.04E-04 | 7.23E-04 | 1.17E-04 | 0.001962 African/African American | 0(v2.1.1), 0(v3.1.2), 0(v4.1.0) | 741416 Likely benign | BS1, BS2 (P), BP4 | Likely Benign |
| CACNA1G | NM_018896.5 | chr17(GRCh38):g.50578473G>A | c.2210G>A | p.Arg737Gln | 2.60E-04 | 4.34E-04 | 1.05E-04 | 0.001095 African/African American | 1(v2.1.1), 0(v3.1.2), 1(v4.1.0) | 2788732 Benign | BS1, BS2 (S), BP4 | Benign |
| CACNA1G | NM_018896.5 | chr17(GRCh38):g.50599434G>A | c.3265G>A | p.Ala1089Thr | 3.46E-04 | 4.99E-04 | 6.77E-04 | 0.0007999 European (non-Finnish) | 0(v2.1.1), 0(v3.1.2), 1(v4.1.0) | 1711441 Likely benign/VUS | BS1, BS2 (P), BP4 | Likely Benign |
| CACNA1G | NM_018896.5 | chr17(GRCh38):g.50599464G>A | c.3295G>A | p.Ala1099Thr | 1.32E-03 | 4.07E-03 | 1.03E-03 | 0.01338 African/African American | 3(v2.1.1), 4(v3.1.2), 14(v4.1.0) | 711189 Benign/Likely benign | BS1, BS2 (S), BP4 | Benign |
| CACNA1G | NM_018896.5 | chr17(GRCh38):g.50600739G>A | c.3704G>A | p.Arg1235Gln | 1.47E-03 | 1.83E-03 | 2.80E-03 | 0.003509 European (non-Finnish) | 1(v2.1.1), 0(v3.1.2), 5(v4.1.0) | 787647 Benign/Likely benign | BS1, BS2 (S), PP3 (P) | Benign |
| CACNA1G | NM_018896.5 | chr17(GRCh38):g.50619003C>T | c.5776C>T | p.Pro1926Ser | 1.60E-04 | 3.94E-04 | 5.78E-05 | 0.0007960 African/African American | 0(v2.1.1), 0(v3.1.2), 0(v4.1.0) | 2049186 Benign/Lilely benign/VUS | BS1, BS2 (P), BP4 | Likely Benign |
| CACNA1G | NM_018896.5 | chr17(GRCh38):g.50624428C>T | c.6298C>T | p.Pro2100Ser | 6.31E-04 | 8.75E-04 | 6.82E-04 | 0.0007339 European (non-Finnish) | 0(v2.1.1), 0(v3.1.2), 2(v4.1.0) | 745634 Likely benign | BS1, BS2 (P), BP4 | Likely Benign |
| CACNA1G | NM_018896.5 | chr17(GRCh38):g.50624452G>T | c.6322G>T | p.Ala2108Ser | 9.21E-03 | 2.72E-02 | 5.40E-03 | 0.09385 African/African American | 94(v2.1.1), 178(v3.1.2), 353(v4.1.0) | 1232827 Benign | BA1, BS2 (S), BP4 | Benign |
| CACNA1G | NM_018896.5 | chr17(GRCh38):g.50626327C>A | c.6710C>A | p.Pro2237His | 4.94E-04 | 4.60E-04 | 4.30E-04 | 0.01540 Middle Eastern | 0(v2.1.1), 0(v3.1.2), 4(v4.1.0) | 279724 Likely benign/VUS | BS1, BS2 (P), BP4 | Likely Benign |
| CACNA1G | NM_018896.5 | chr17(GRCh38):g.50626582G>A | c.6965G>A | p.Arg2322Gln | 5.70E-04 | 1.63E-03 | 3.01E-04 | 0.005313 African/African American | 1(v2.1.1),0(v3.1.2), 2(v4.1.0) | 711659 Benign | BS1, BS2 (S), BP4 (S) | Benign |
| CACNA1G | NM_018896.5 | chr17(GRCh38):g.50626594G>A | c.6977G>A | p.Ser2326Asn | 4.78E-04 | 1.40E-04 | 2.27E-04 | 0.003404 South Asian | 0(v2.1.1), 0(v3.1.2), 1(v4.1.0) | 3046754 Likely benign | BS1, BS2 (P), BP4 (S) | Benign |
| CACNA1G | NM_018896.5 | chr17(GRCh38):g.50626683C>T | c.7066C>T | p.Pro2356Ser | 3.32E-04 | 4.60E-04 | 4.62E-04 | 0.0005483 European (Finnish) | 0(v2.1.1), 0(v3.1.2), 0(v4.1.0) | 374702 Likely benign | BS1, BS2 (P), BP4 | Likely Benign |

**Pathogenic controls**

| Gene | Transcript | Genomic Location and Change | DNA Change | Protein Change | gnomAD AF | | | gnomAD v4.1.0 Grpmax Filtering AF | gnomAD homozygotes | Clinvar Classification (19 Jan 26) | ACMG/AMP Criteria | Final Classification |
|---|---|---|---|---|---|---|---|---|---|---|---|---|
| | | | | | V2.1.1 (Exomes) | v3.1.2 | v4.1.0 (Total) | | | | | |
| CACNA1G | NM_018896.5 | chr17(GRCh38):g.50569291A>T | c.481A>T | p.Ile161Phe | 1.20E-05 | 7.05E-06 | 2.48E-06 | N/A | 0(2.1.1),0(3.1.2), 0(v4.1.0) | 1027470 Likely pathogenic/VUS | PM2(P), PM6(M), PP2, PP3(M), | Likely Pathogenic |
| CACNA1G | NM_018896.5 | chr17(GRCh38):g.50571881T>G | c.590T>G | p.Met197Arg | - | - | - | | - | 1722930 Likely pathogenic | PS2(S), PM2(P), PP2, PP3(S) | Pathogenic |
| CACNA1G | NM_018896.5 | chr17(GRCh38):g.50571914T>C | c.623T>C | p.Leu208Pro | - | - | - | | - | 452482 Likely pathogenic | PS2 (S), PM2(P), PP2, PP3(S) | Pathogenic |
| CACNA1G | NM_018896.5 | chr17(GRCh38):g.50571923T>C | c.632T>C | p.Leu211Pro | - | - | - | | - | 974856 Likely pathogenic | PM2 (P), PM6(M), PP2, PP3(S) | Likely Pathogenic |
| CACNA1G | NM_018896.5 | chr17(GRCh38):g.50573076C>T | c.1103C>T | p.Ser368Phe | - | - | - | | - | 871462 Likely pathogenic | PM2 (P), PP2, PP3(S) | Likely Pathogenic |
| CACNA1G | NM_018896.5 | chr17(GRCh38):g.50591580A>C | c.2599A>C | p.Thr867Pro | - | - | - | | - | 265067 Likely pathogenic | PM2 (P), PP2, PP3(S) | Likely Pathogenic |
| CACNA1G | NM_018896.5 | chr17(GRCh38):g.50591992C>T | c.2810C>T | p.Ser937Leu | 4.02E-06 | - | 1.24E-06 | N/A | 0(2.1.1), -, 0(v4.1.0) | 1228386 Likely pathogenic | PM2(P), PM6(M), PP2, PP3(S) | Likely Pathogenic |
| CACNA1G | NM_018896.5 | chr17(GRCh38):g.50592063G>A | c.2881G>A | p.Ala961Thr | - | - | - | | - | 280269 Pathogenic | PS2 (VS), PS4(M), PM1, PM2 (P), PP3(S) | Pathogenic |
| CACNA1G | NM_018896.5 | chr17(GRCh38):g.50601094G>A | c.3835G>A | p.Asp1279Asn | - | - | 6.20E-07 | N/A | - | 1027472 Likely pathogenic | PS4 (M), PM2 (P), PM6(M) PP2, PP3(M) | Likely Pathogenic |
| CACNA1G | NM_018896.5 | chr17(GRCh38):g.50607905A>G | c.4591A>G | p.Met1531Val | - | - | - | | - | 559581 Pathogenic | PS2 (VS), PS4(M), PM1, PM2 (P), PP3(S) | Pathogenic |
| CACNA1G | NM_018896.5 | chr17(GRCh38):g.50607906T>C | c.4592T>C | p.Met1531Thr | - | - | - | | - | 1804078 Likely pathogenic | PM2(P), PM1, PP3(S) | Likely Pathogenic |
| CACNA1G | NM_018896.5 | chr17(GRCh38):g.50607908T>C | c.4594T>C | p.Phe1532Leu | - | - | - | | - | 1332877 Likely pathogenic | PM2(P),PM1, PM6(M), PP3(S) | Likely Pathogenic |
| CACNA1G | NM_018896.5 | chr17(GRCh38):g.50607915G>A | c.4601G>A | p.Gly1534Asp | - | - | - | | - | 808297 Likely pathogenic | PS2 (S), PM1, PM2(P), PP3(S) | Pathogenic |
| CACNA1G | NM_018896.5 | chr17(GRCh38):g.50617560G>A | c.5144G>A | p.Arg1715His | - | - | - | | - | 221981 Pathogenic | PS4(M), PM2 (P), PP1 (S), PP2, PP3(S) | Pathogenic |
| CACNA1G | NM_018896.5 | chr17(GRCh38):g.50617568C>G | c.5152C>G | p.Arg1718Gly | - | - | - | | - | 1700076 Likely pathogenic | PM2 (P), PM6(M), PP2, PP3(S) | Likely Pathogenic |
| CACNA1G | NM_018896.5 | chr17(GRCh38):g.50618311T>A | c.5395T>A | p.Ser1799Thr | - | - | - | | - | 1027464 Likely pathogenic | PM2 (P), PM6(M), PP2, PP3(M) | Likely Pathogenic |

**VUS**

| Gene | Transcript | Genomic Location and Change | DNA Change | Protein Change | gnomAD AF | | | gnomAD v4.1.0 Grpmax Filtering AF | gnomAD homozygotes | Clinvar Classification (19 Jan 26) | ACMG/AMP Criteria | Final Classification |
|---|---|---|---|---|---|---|---|---|---|---|---|---|
| | | | | | V2.1.1 (Exomes) | v3.1.2 | v4.1.0 (Total) | | | | | |
| CACNA1G | NM_018896.5 | chr17(GRCh38):g.50569774G>A | c.557G>A | p.Arg186Gln | - | - | 6.45E-07 | N/A | -, -, 0(v4.1.0) | 4542197 VUS | PM2, PM1, PP3, PS3 | Likely Pathogenic |
| CACNA1G | NM_018896.5 | chr17(GRCh38):g.50599462G>A | c.3293G>A | p.Ser1098Asn | 9.29E-06 | 1.31E-05 | 1.08E-05 | 0.000008920 European (non-Finnish) | 0(v2.1.1), 0(v3.1.2), 0(v4.1.0) | NA | PM2 (P), PP2, BP4 | VUS |
| CACNA1G | NM_018896.5 | chr17(GRCh38):g.50604166G>A | c.4181G>A | p.Arg1394Gln | - | - | 2.48E-06 | 0.00000068 European (non-Finnish) | -, -, 0(v4.1.0) | 3773799 VUS | PM2, PP2, PP3, PS3 | Likely Pathogenic |
| CACNA1G | NM_018896.5 | chr17(GRCh38):g.50624506G>T | c.6376G>T | p.Ala2126Ser | - | - | 3.13E-06 | 0.000001250 European (non-Finnish) | -, -, 0(v4.1.0) | 1706111 VUS | PM2 (P), PP2, BP4 | VUS |
| CACNA1G | NM_018896.5 | chr17(GRCh38):g.50626063G>A | c.6446G>A | p.Arg2149Gln | 2.43E-05 | 1.97E-05 | 3.72E-05 | 0.00003585 European (non-Finnish) | 0(v2.1.1), 0(v3.1.2), 0(v4.1.0) | 2897851 Likely benign | PM2 (P), PP2, BP4 | VUS |
