## Supplemental Table 2 for "Calibrated high-throughput electrophysiology enables clinical interpretation of *CACNA1G* missense variants"

**Table S2: Power calculation**

| | assuming K=1 | | | | | Sample size |
|---|---|---|---|---|---|---|
| **CD*** | **Cav3.1WT** | | | | | |
| | 33.78 | n1 | 1140.78 | 2281.56 | 10.50 | n |
| | | | | 25 | 625 | 38 |
| | | | | 50 | 2500 | 10 |
| **$\tau_{Deact}$'** | **Cav3.1WT** | | | | | |
| | 22.42 | n1 | 502.66 | 1005.31 | 10.50 | n |
| | | | | 25 | 625 | 17 |
| | | | | 50 | 2500 | 4 |
