## Supplemental Table 3 for "Calibrated high-throughput electrophysiology enables clinical interpretation of *CACNA1G* missense variants"

**Table S3: Square-root transformed and WT normalized current density**

| Column | Mean | StdDev | N | 95% CI Lower | 95% CI Upper | Z Score | Benign data |
|---|---|---|---|---|---|---|---|
| Cav3.1_WT | 1.00 | 0.34 | 1179 | 0.02 | 0.02 | -0.39 | |
| Cav3.1_V96M | 0.93 | 0.33 | 237 | 0.04 | 0.04 | -1.06 | mean |
| Cav3.1_I161F | 1.06 | 0.44 | 194 | 0.06 | 0.06 | 0.20 | 1.04 |
| Cav3.1_R186Q | 0.67 | 0.24 | 174 | 0.04 | 0.04 | -3.61 | std_dev |
| Cav3.1_M197R | 0.37 | 0.12 | 125 | 0.02 | 0.02 | -6.64 | 0.10 |
| Cav3.1_L208P | 0.51 | 0.17 | 140 | 0.03 | 0.03 | -5.18 | n |
| Cav3.1_L211P | 0.43 | 0.21 | 276 | 0.02 | 0.02 | -5.97 | 25 |
| Cav3.1_G299S | 1.17 | 0.47 | 214 | 0.06 | 0.06 | 1.30 | |
| Cav3.1_S368F | 0.58 | 0.22 | 272 | 0.03 | 0.03 | -4.50 | |
| Cav3.1_R456Q | 0.98 | 0.39 | 100 | 0.08 | 0.08 | -0.51 | |
| Cav3.1_R530H | 1.13 | 0.44 | 167 | 0.07 | 0.07 | 0.93 | |
| Cav3.1_T539M | 1.13 | 0.40 | 110 | 0.08 | 0.08 | 0.96 | |
| Cav3.1_A545P | 0.96 | 0.31 | 120 | 0.06 | 0.06 | -0.77 | |
| Cav3.1_G549A | 1.22 | 0.48 | 191 | 0.07 | 0.07 | 1.80 | |
| Cav3.1_A567C | 1.10 | 0.40 | 274 | 0.05 | 0.05 | 0.67 | |
| Cav3.1_V593M | 1.08 | 0.39 | 126 | 0.07 | 0.07 | 0.42 | |
| Cav3.1_G627R | 1.07 | 0.41 | 129 | 0.07 | 0.07 | 0.31 | |
| Cav3.1_C644Y | 0.97 | 0.36 | 113 | 0.07 | 0.07 | -0.62 | |
| Cav3.1_S690N | 1.11 | 0.41 | 150 | 0.07 | 0.07 | 0.76 | |
| Cav3.1_R714W | 0.97 | 0.38 | 243 | 0.05 | 0.05 | -0.67 | |
| Cav3.1_S715I | 1.19 | 0.41 | 222 | 0.05 | 0.05 | 1.51 | |
| Cav3.1_R737Q | 1.09 | 0.36 | 164 | 0.06 | 0.06 | 0.54 | |
| Cav3.1_T867P | 0.88 | 0.34 | 134 | 0.06 | 0.06 | -1.51 | |
| Cav3.1_S937L | 0.52 | 0.18 | 138 | 0.03 | 0.03 | -5.10 | |
| Cav3.1_A961T | 0.62 | 0.31 | 110 | 0.06 | 0.06 | -4.14 | |
| Cav3.1_A1089T | 0.80 | 0.31 | 144 | 0.05 | 0.05 | -2.34 | |
| Cav3.1_S1098N | 1.07 | 0.37 | 159 | 0.06 | 0.06 | 0.33 | |
| Cav3.1_A1099T | 0.94 | 0.33 | 129 | 0.06 | 0.06 | -0.90 | |
| Cav3.1_R1235Q | 0.94 | 0.41 | 93 | 0.08 | 0.08 | -0.91 | |
| Cav3.1_D1279N | 0.81 | 0.28 | 125 | 0.05 | 0.05 | -2.26 | |
| Cav3.1_R1394Q | 1.12 | 0.42 | 79 | 0.09 | 0.09 | 0.82 | |
| Cav3.1_M1531T | 0.64 | 0.22 | 108 | 0.04 | 0.04 | -3.91 | |
| Cav3.1_M1531V | 0.64 | 0.24 | 136 | 0.04 | 0.04 | -3.91 | |
| Cav3.1_F1532L | 0.69 | 0.21 | 124 | 0.04 | 0.04 | -3.39 | |
| Cav3.1_G1534D | 0.33 | 0.13 | 126 | 0.02 | 0.02 | -7.05 | |
| Cav3.1_R1715H | 0.83 | 0.31 | 129 | 0.05 | 0.05 | -2.08 | |
| Cav3.1_R1718G | 0.69 | 0.22 | 219 | 0.03 | 0.03 | -3.48 | |
| Cav3.1_S1799T | 0.99 | 0.40 | 127 | 0.07 | 0.07 | -0.45 | |
| Cav3.1_P1926S | 1.06 | 0.38 | 104 | 0.07 | 0.07 | 0.25 | |
| Cav3.1_P2100S | 1.07 | 0.33 | 128 | 0.06 | 0.06 | 0.32 | |
| Cav3.1_A2108S | 1.07 | 0.45 | 93 | 0.09 | 0.09 | 0.38 | |
| Cav3.1_A2126S | 1.01 | 0.40 | 99 | 0.08 | 0.08 | -0.20 | |
| Cav3.1_R2149Q | 1.00 | 0.36 | 128 | 0.06 | 0.06 | -0.39 | |
| Cav3.1_P2237H | 1.05 | 0.48 | 122 | 0.09 | 0.09 | 0.16 | |
| Cav3.1_S2326N | 0.93 | 0.51 | 129 | 0.09 | 0.09 | -1.09 | |
| Cav3.1_R2322Q | 1.03 | 0.48 | 165 | 0.07 | 0.07 | -0.06 | |
| Cav3.1_P2356S | 0.90 | 0.46 | 83 | 0.10 | 0.10 | -1.39 | |
| NEG | 0.40 | 0.18 | 806 | 0.01 | 0.01 | -6.32 | |
