## Supplemental Table 4 for "Calibrated high-throughput electrophysiology enables clinical interpretation of *CACNA1G* missense variants"

**Table S4: Calibration of functional evidence strength**

| Dataset | A | B | C | D | P1 | P2path | P2benign | OddsPath_P | OddsPath_B | Functionally Abnormal Code | Functionally Normal Code |
|---|---|---|---|---|---|---|---|---|---|---|---|
| CACNA1G CD* | 24 | 1 | 3 | 13 | 0.390 | 0.929 | 0.111 | 20.3 | 0.20 | PS3_strong | BS3_moderate* |
| CACNA1G tau Deact' | 24 | 1 | 2 | 11 | 0.342 | 0.917 | 0.077 | 21.2 | 0.16 | PS3_strong | BS3_supporting |

A:  Benign + Normal assay result
B: Benign and Abnormal assay result
C: Pathogenic and Normal assay result
D: Pathogenic and Abnormal assay result
