## Supplemental Table 5 for "Calibrated high-throughput electrophysiology enables clinical interpretation of *CACNA1G* missense variants"

**Table S5: Summary of electrophysiological data for channel activation**

| Column | Mean | StdDev | N | 95% CI Lower | 95% CI Upper | Z Score | Benign data |
|---|---|---|---|---|---|---|---|
| Cav3.1_WT | 0.29 | 5.11 | 784 | 0.36 | 0.36 | 0.24 | |
| Cav3.1_V96M | 1.85 | 6.34 | 129 | 1.11 | 1.11 | 0.97 | mean |
| Cav3.1_I161F | 2.21 | 6.71 | 119 | 1.22 | 1.22 | 1.15 | -0.21 |
| Cav3.1_R186Q | -5.80 | 5.91 | 104 | 1.15 | 1.15 | -2.65 | std_dev |
| Cav3.1_M197R | -- | -- | -- | -- | -- | -- | 2.11 |
| Cav3.1_L208P | -0.56 | 6.81 | 27 | 2.69 | 2.69 | -0.16 | n |
| Cav3.1_L211P | -- | -- | -- | -- | -- | -- | 25 |
| Cav3.1_G299S | 0.97 | 6.40 | 137 | 1.08 | 1.08 | 0.56 | |
| Cav3.1_S368F | 2.82 | 4.93 | 67 | 1.20 | 1.20 | 1.44 | |
| Cav3.1_R456Q | -0.47 | 4.96 | 69 | 1.19 | 1.19 | -0.12 | |
| Cav3.1_R530H | 2.20 | 6.45 | 112 | 1.21 | 1.21 | 1.14 | |
| Cav3.1_T539M | -1.41 | 5.18 | 88 | 1.10 | 1.10 | -0.57 | |
| Cav3.1_A545P | -2.82 | 5.96 | 77 | 1.35 | 1.35 | -1.23 | |
| Cav3.1_G549A | 2.03 | 6.59 | 131 | 1.14 | 1.14 | 1.06 | |
| Cav3.1_A567C | 1.00 | 6.42 | 185 | 0.93 | 0.93 | 0.57 | |
| Cav3.1_V593M | 0.11 | 7.22 | 91 | 1.50 | 1.50 | 0.15 | |
| Cav3.1_G627R | -1.09 | 4.75 | 76 | 1.08 | 1.08 | -0.41 | |
| Cav3.1_C644Y | -4.19 | 5.71 | 65 | 1.41 | 1.41 | -1.88 | |
| Cav3.1_S690N | -0.91 | 4.89 | 90 | 1.02 | 1.02 | -0.33 | |
| Cav3.1_R714W | -1.95 | 6.19 | 132 | 1.07 | 1.07 | -0.82 | |
| Cav3.1_S715I | 2.10 | 6.20 | 157 | 0.98 | 0.98 | 1.09 | |
| Cav3.1_R737Q | -1.11 | 5.98 | 118 | 1.09 | 1.09 | -0.42 | |
| Cav3.1_T867P | -12.00 | 4.32 | 63 | 1.09 | 1.09 | -5.58 | |
| Cav3.1_S937L | 3.78 | 5.78 | 11 | 3.89 | 3.89 | 1.89 | |
| Cav3.1_A961T | -1.77 | 6.78 | 32 | 2.44 | 2.44 | -0.74 | |
| Cav3.1_A1089T | -2.03 | 5.15 | 97 | 1.04 | 1.04 | -0.86 | |
| Cav3.1_S1098N | 0.53 | 5.16 | 110 | 0.97 | 0.97 | 0.35 | |
| Cav3.1_A1099T | -3.53 | 5.34 | 89 | 1.12 | 1.12 | -1.57 | |
| Cav3.1_R1235Q | -3.98 | 8.16 | 55 | 2.21 | 2.21 | -1.79 | |
| Cav3.1_D1279N | 1.36 | 5.61 | 60 | 1.45 | 1.45 | 0.74 | |
| Cav3.1_R1394Q | 5.75 | 5.06 | 53 | 1.40 | 1.40 | 2.82 | |
| Cav3.1_M1531T | -4.21 | 4.76 | 48 | 1.38 | 1.38 | -1.89 | |
| Cav3.1_M1531V | -6.89 | 5.26 | 34 | 1.84 | 1.84 | -3.16 | |
| Cav3.1_F1532L | -9.61 | 6.27 | 67 | 1.53 | 1.53 | -4.45 | |
| Cav3.1_G1534D | -- | -- | -- | -- | -- | -- | |
| Cav3.1_R1715H | 6.11 | 5.86 | 84 | 1.27 | 1.27 | 2.99 | |
| Cav3.1_R1718G | 4.23 | 4.37 | 136 | 0.74 | 0.74 | 2.10 | |
| Cav3.1_S1799T | -7.86 | 6.89 | 81 | 1.52 | 1.52 | -3.62 | |
| Cav3.1_P1926S | 2.60 | 4.69 | 74 | 1.09 | 1.09 | 1.33 | |
| Cav3.1_P2100S | -0.25 | 5.87 | 99 | 1.17 | 1.17 | -0.02 | |
| Cav3.1_A2108S | 0.72 | 9.95 | 43 | 3.06 | 3.06 | 0.44 | |
| Cav3.1_A2126S | 0.98 | 5.07 | 71 | 1.20 | 1.20 | 0.57 | |
| Cav3.1_R2149Q | 3.22 | 6.19 | 96 | 1.25 | 1.25 | 1.62 | |
| Cav3.1_P2237H | 3.67 | 8.42 | 67 | 2.05 | 2.05 | 1.84 | |
| Cav3.1_S2326N | 0.81 | 7.83 | 66 | 1.92 | 1.92 | 0.48 | |
| Cav3.1_R2322Q | -0.45 | 7.11 | 74 | 1.65 | 1.65 | -0.11 | |
| Cav3.1_P2356S | 0.82 | 11.23 | 36 | 3.80 | 3.80 | 0.49 | |
| Cav3.1_NEG | -- | -- | -- | -- | -- | -- | |
