## Supplemental Table 6 for "Calibrated high-throughput electrophysiology enables clinical interpretation of *CACNA1G* missense variants"

**Table S6: Summary of electrophysiological data for channel inactivation**

| Column | Mean | StdDev | N | 95% CI Lower | 95% CI Upper | Z Score | Benign data |
|---|---|---|---|---|---|---|---|
| Cav3.1_WT | 0.08 | 3.94 | 899 | 0.26 | 0.26 | 0.19 | |
| Cav3.1_V96M | 0.68 | 4.86 | 175 | 0.73 | 0.73 | 0.55 | mean |
| Cav3.1_I161F | 1.22 | 5.67 | 147 | 0.92 | 0.92 | 0.86 | -0.24 |
| Cav3.1_R186Q | -7.52 | 3.79 | 148 | 0.62 | 0.62 | -4.30 | std_dev |
| Cav3.1_M197R | -- | -- | -- | -- | -- | -- | 1.69 |
| Cav3.1_L208P | -20.83 | 11.51 | 3 | 28.60 | 28.60 | -12.17 | n |
| Cav3.1_L211P | -- | -- | -- | -- | -- | -- | 25 |
| Cav3.1_G299S | 1.48 | 3.99 | 172 | 0.60 | 0.60 | 1.02 | |
| Cav3.1_S368F | -2.24 | 4.43 | 65 | 1.10 | 1.10 | -1.18 | |
| Cav3.1_R456Q | -0.12 | 3.42 | 78 | 0.77 | 0.77 | 0.07 | |
| Cav3.1_R530H | 1.75 | 4.58 | 135 | 0.78 | 0.78 | 1.18 | |
| Cav3.1_T539M | -0.35 | 3.79 | 96 | 0.77 | 0.77 | -0.06 | |
| Cav3.1_A545P | -3.19 | 4.44 | 87 | 0.95 | 0.95 | -1.74 | |
| Cav3.1_G549A | 1.79 | 3.90 | 152 | 0.62 | 0.62 | 1.20 | |
| Cav3.1_A567C | 1.08 | 4.13 | 207 | 0.57 | 0.57 | 0.78 | |
| Cav3.1_V593M | 0.64 | 4.64 | 98 | 0.93 | 0.93 | 0.53 | |
| Cav3.1_G627R | -0.29 | 3.91 | 92 | 0.81 | 0.81 | -0.03 | |
| Cav3.1_C644Y | -3.82 | 4.89 | 78 | 1.10 | 1.10 | -2.11 | |
| Cav3.1_S690N | -1.24 | 3.74 | 102 | 0.74 | 0.74 | -0.59 | |
| Cav3.1_R714W | -0.59 | 5.52 | 177 | 0.82 | 0.82 | -0.21 | |
| Cav3.1_S715I | 1.80 | 4.45 | 179 | 0.66 | 0.66 | 1.21 | |
| Cav3.1_R737Q | -0.02 | 4.50 | 128 | 0.79 | 0.79 | 0.14 | |
| Cav3.1_T867P | -6.12 | 5.91 | 112 | 1.11 | 1.11 | -3.47 | |
| Cav3.1_S937L | 0.72 | 3.16 | 10 | 2.26 | 2.26 | 0.57 | |
| Cav3.1_A961T | -2.80 | 5.23 | 29 | 1.99 | 1.99 | -1.51 | |
| Cav3.1_A1089T | -2.02 | 3.52 | 102 | 0.69 | 0.69 | -1.05 | |
| Cav3.1_S1098N | 0.67 | 3.34 | 138 | 0.56 | 0.56 | 0.54 | |
| Cav3.1_A1099T | -2.75 | 4.06 | 95 | 0.83 | 0.83 | -1.48 | |
| Cav3.1_R1235Q | -2.28 | 6.23 | 69 | 1.50 | 1.50 | -1.20 | |
| Cav3.1_D1279N | -1.88 | 4.60 | 71 | 1.09 | 1.09 | -0.97 | |
| Cav3.1_R1394Q | 3.96 | 3.32 | 61 | 0.85 | 0.85 | 2.49 | |
| Cav3.1_M1531T | -2.21 | 4.32 | 66 | 1.06 | 1.06 | -1.16 | |
| Cav3.1_M1531V | -5.75 | 4.61 | 44 | 1.40 | 1.40 | -3.26 | |
| Cav3.1_F1532L | -7.50 | 4.61 | 89 | 0.97 | 0.97 | -4.29 | |
| Cav3.1_G1534D | -- | -- | -- | -- | -- | -- | |
| Cav3.1_R1715H | -1.65 | 4.11 | 84 | 0.89 | 0.89 | -0.83 | |
| Cav3.1_R1718G | -3.04 | 3.30 | 141 | 0.55 | 0.55 | -1.65 | |
| Cav3.1_S1799T | -2.29 | 5.26 | 107 | 1.01 | 1.01 | -1.21 | |
| Cav3.1_P1926S | 2.26 | 3.87 | 82 | 0.85 | 0.85 | 1.48 | |
| Cav3.1_P2100S | 0.42 | 4.33 | 102 | 0.85 | 0.85 | 0.39 | |
| Cav3.1_A2108S | -0.32 | 7.58 | 50 | 2.15 | 2.15 | -0.04 | |
| Cav3.1_A2126S | 0.87 | 3.22 | 82 | 0.71 | 0.71 | 0.66 | |
| Cav3.1_R2149Q | 1.63 | 3.66 | 107 | 0.70 | 0.70 | 1.11 | |
| Cav3.1_P2237H | 2.05 | 5.68 | 75 | 1.31 | 1.31 | 1.35 | |
| Cav3.1_S2326N | -1.13 | 8.03 | 85 | 1.73 | 1.73 | -0.52 | |
| Cav3.1_R2322Q | -1.04 | 5.70 | 97 | 1.15 | 1.15 | -0.47 | |
| Cav3.1_P2356S | -0.89 | 7.85 | 56 | 2.10 | 2.10 | -0.38 | |
| Cav3.1_NEG | -- | -- | -- | -- | -- | -- | |
