## Supplemental Table 7 for "Calibrated high-throughput electrophysiology enables clinical interpretation of *CACNA1G* missense variants"

**Table S7: Summary of electrophysiological data for channel deactivation**

| Column | Mean | StdDev | N | 95% CI Lower | 95% CI Upper | Z Score | Benign data |
|---|---|---|---|---|---|---|---|
| Cav3.1_WT | 0.41 | 0.09 | 558 | 0.01 | 0.01 | -1.04 | |
| Cav3.1_V96M | 0.38 | 0.10 | 100 | 0.02 | 0.02 | -2.00 | mean |
| Cav3.1_I161F | 0.46 | 0.10 | 73 | 0.02 | 0.02 | 0.18 | 0.45 |
| Cav3.1_R186Q | 0.67 | 0.11 | 120 | 0.02 | 0.02 | 6.31 | std_dev |
| Cav3.1_M197R | -- | -- | -- | -- | -- | -- | 0.04 |
| Cav3.1_L208P | 0.91 | 0.14 | 32 | 0.05 | 0.05 | 13.04 | n |
| Cav3.1_L211P | -- | -- | -- | -- | -- | -- | 25 |
| Cav3.1_G299S | 0.46 | 0.09 | 95 | 0.02 | 0.02 | 0.31 | |
| Cav3.1_S368F | 0.35 | 0.08 | 58 | 0.02 | 0.02 | -2.72 | |
| Cav3.1_R456Q | 0.51 | 0.12 | 55 | 0.03 | 0.03 | 1.73 | |
| Cav3.1_R530H | 0.42 | 0.09 | 56 | 0.02 | 0.02 | -0.94 | |
| Cav3.1_T539M | 0.45 | 0.08 | 43 | 0.02 | 0.02 | -0.01 | |
| Cav3.1_A545P | 0.45 | 0.07 | 50 | 0.02 | 0.02 | 0.12 | |
| Cav3.1_G549A | 0.43 | 0.08 | 66 | 0.02 | 0.02 | -0.62 | |
| Cav3.1_A567C | 0.44 | 0.09 | 114 | 0.02 | 0.02 | -0.31 | |
| Cav3.1_V593M | 0.51 | 0.13 | 34 | 0.04 | 0.04 | 1.66 | |
| Cav3.1_G627R | 0.46 | 0.11 | 39 | 0.04 | 0.04 | 0.28 | |
| Cav3.1_C644Y | 0.47 | 0.07 | 37 | 0.02 | 0.02 | 0.59 | |
| Cav3.1_S690N | 0.42 | 0.08 | 55 | 0.02 | 0.02 | -0.85 | |
| Cav3.1_R714W | 0.40 | 0.09 | 106 | 0.02 | 0.02 | -1.46 | |
| Cav3.1_S715I | 0.49 | 0.10 | 102 | 0.02 | 0.02 | 1.12 | |
| Cav3.1_R737Q | 0.49 | 0.10 | 31 | 0.04 | 0.04 | 1.20 | |
| Cav3.1_T867P | 0.64 | 0.11 | 40 | 0.04 | 0.04 | 5.39 | |
| Cav3.1_S937L | 0.46 | 0.15 | 7 | 0.14 | 0.14 | 0.26 | |
| Cav3.1_A961T | 0.79 | 0.09 | 24 | 0.04 | 0.04 | 9.83 | |
| Cav3.1_A1089T | 0.46 | 0.07 | 54 | 0.02 | 0.02 | 0.26 | |
| Cav3.1_S1098N | 0.48 | 0.10 | 98 | 0.02 | 0.02 | 0.92 | |
| Cav3.1_A1099T | 0.43 | 0.07 | 55 | 0.02 | 0.02 | -0.47 | |
| Cav3.1_R1235Q | 0.47 | 0.08 | 15 | 0.05 | 0.05 | 0.70 | |
| Cav3.1_D1279N | 0.60 | 0.08 | 54 | 0.02 | 0.02 | 4.21 | |
| Cav3.1_R1394Q | 0.09 | 0.09 | 33 | 0.03 | 0.03 | -10.17 | |
| Cav3.1_M1531T | 0.52 | 0.07 | 29 | 0.02 | 0.02 | 2.08 | |
| Cav3.1_M1531V | 0.88 | 0.14 | 71 | 0.03 | 0.03 | 12.34 | |
| Cav3.1_F1532L | 0.73 | 0.13 | 28 | 0.05 | 0.05 | 7.98 | |
| Cav3.1_G1534D | -- | -- | -- | -- | -- | -- | |
| Cav3.1_R1715H | 0.27 | 0.09 | 53 | 0.02 | 0.02 | -5.01 | |
| Cav3.1_R1718G | 0.35 | 0.08 | 140 | 0.01 | 0.01 | -2.73 | |
| Cav3.1_S1799T | 0.74 | 0.09 | 28 | 0.04 | 0.04 | 8.21 | |
| Cav3.1_P1926S | 0.47 | 0.10 | 35 | 0.03 | 0.03 | 0.52 | |
| Cav3.1_P2100S | 0.45 | 0.09 | 52 | 0.02 | 0.02 | 0.05 | |
| Cav3.1_A2108S | 0.47 | 0.09 | 13 | 0.05 | 0.05 | 0.58 | |
| Cav3.1_A2126S | 0.50 | 0.12 | 66 | 0.03 | 0.03 | 1.43 | |
| Cav3.1_R2149Q | 0.48 | 0.09 | 92 | 0.02 | 0.02 | 0.87 | |
| Cav3.1_P2237H | 0.40 | 0.10 | 57 | 0.03 | 0.03 | -1.48 | |
| Cav3.1_S2326N | 0.49 | 0.12 | 61 | 0.03 | 0.03 | 1.06 | |
| Cav3.1_R2322Q | 0.41 | 0.10 | 65 | 0.02 | 0.02 | -1.09 | |
| Cav3.1_P2356S | 0.42 | 0.10 | 33 | 0.04 | 0.04 | -0.93 | |
| NEG | -- | -- | -- | -- | -- | -- | |
