## Supplemental Figure 1 for "Calibrated high-throughput electrophysiology enables clinical interpretation of *CACNA1G* missense variants"

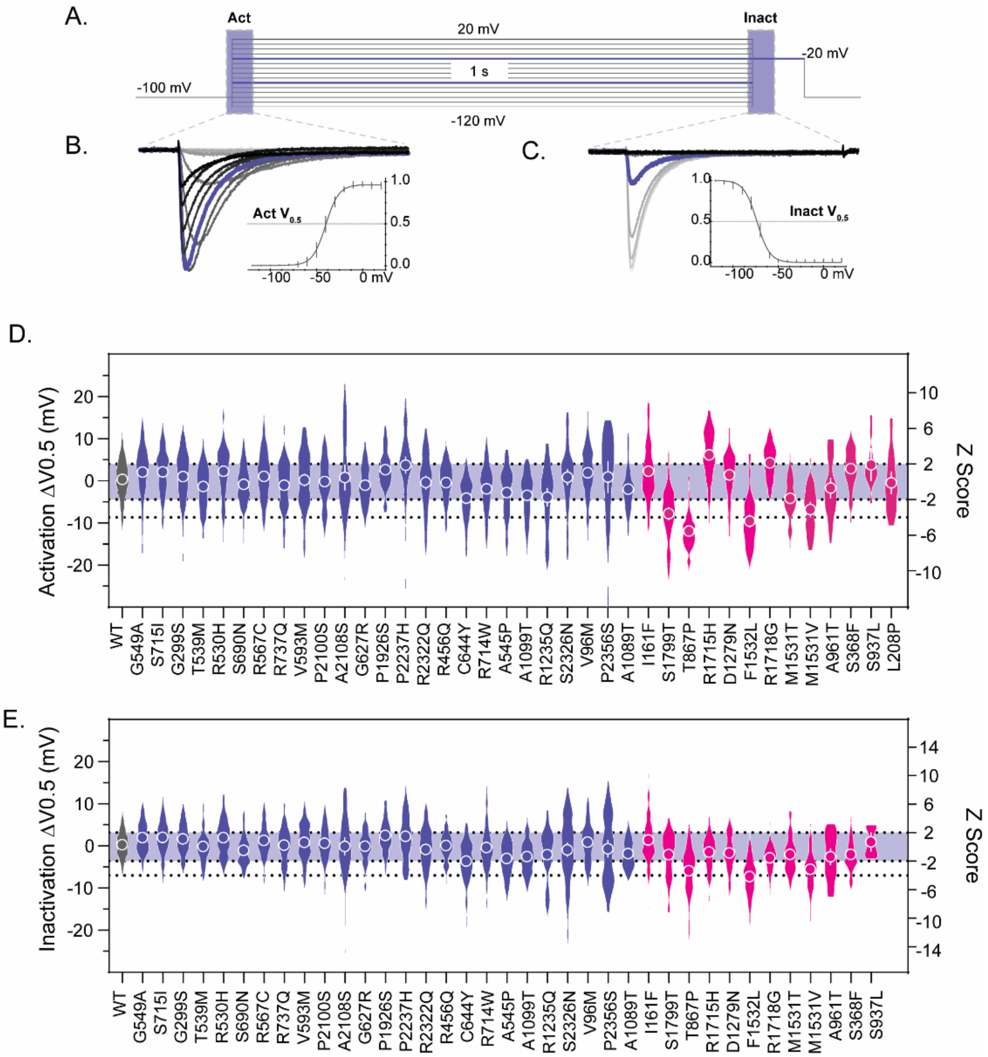


**Figure S1**. **Evaluation of voltage-dependence provides limited but supportive functional context for *CACNA1G* variant interpretation. A.** Voltage-clamp protocol used to assess activation and inactivation, employing depolarizing steps from –120 mV to +20 mV (Vh: –100 mV, 0.1 Hz). **B–C**. Representative WT current traces used to generate activation plots (peak inward currents during first 25 ms) and inactivation plots (relative current measured at –20 mV after 1 s depolarizing pulses). Relative voltage dependence was quantified as the difference in half-activation (act $\Delta V_{0.5}$) and half-inactivation (inact $\Delta V_{0.5}$) potentials between each variant and WT. **D–E**. Mean ± 95% CI for act $\Delta V_{0.5}$ and inact $\Delta V_{0.5}$, shown with raw values (left axis) and corresponding Z-scores (right axis). Normal functional range (mean ± 2 SD of benign values) is shaded in blue; dotted lines indicate values beyond ±2. B/LB variants are shown in blue and P/LP variants in magenta. Variant-specific sample sizes are provided in Table S5.
