## Supplemental Figure 2 for "Calibrated high-throughput electrophysiology enables clinical interpretation of *CACNA1G* missense variants"

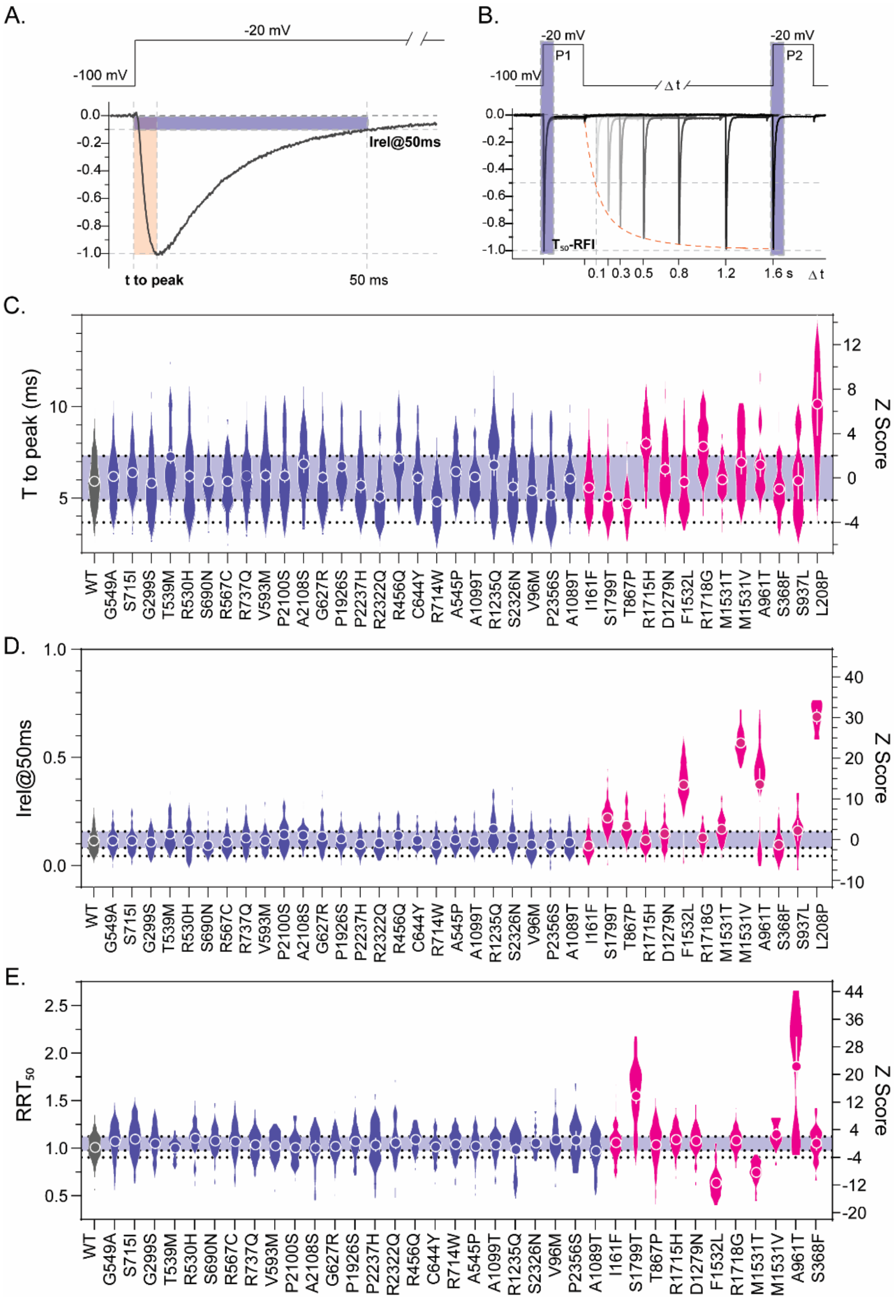


**Figure S2**. **Evaluation of current-kinetic parameters provides supportive functional context for *CACNA1G* variant interpretation**. **A.** Representative WT current traces and voltage-clamp protocols used to assess activation kinetics (time to peak), inactivation kinetics (Irel@50ms), and recovery from inactivation (Relative Recovery Time, $RRT_{50}$). $RRT_{50}$ is defined as the ratio of the time required for 50% recovery in a variant relative to WT. Panels **C–E** show mean ± 95% CI for time to peak, Irel@50ms, and $RRT_{50}$, shown with corresponding raw values (left axis) and corresponding Z-scores (right axis). Normal functional range (mean ± 2 SD of benign values) is shaded in blue; dotted lines indicate values beyond ±2. B/LB variants are shown in blue and P/LP variants in magenta. Variant-specific sample sizes are provided in Table S8, S9 and S9.
