## Supplemental Figure 3 for "Calibrated high-throughput electrophysiology enables clinical interpretation of *CACNA1G* missense variants"

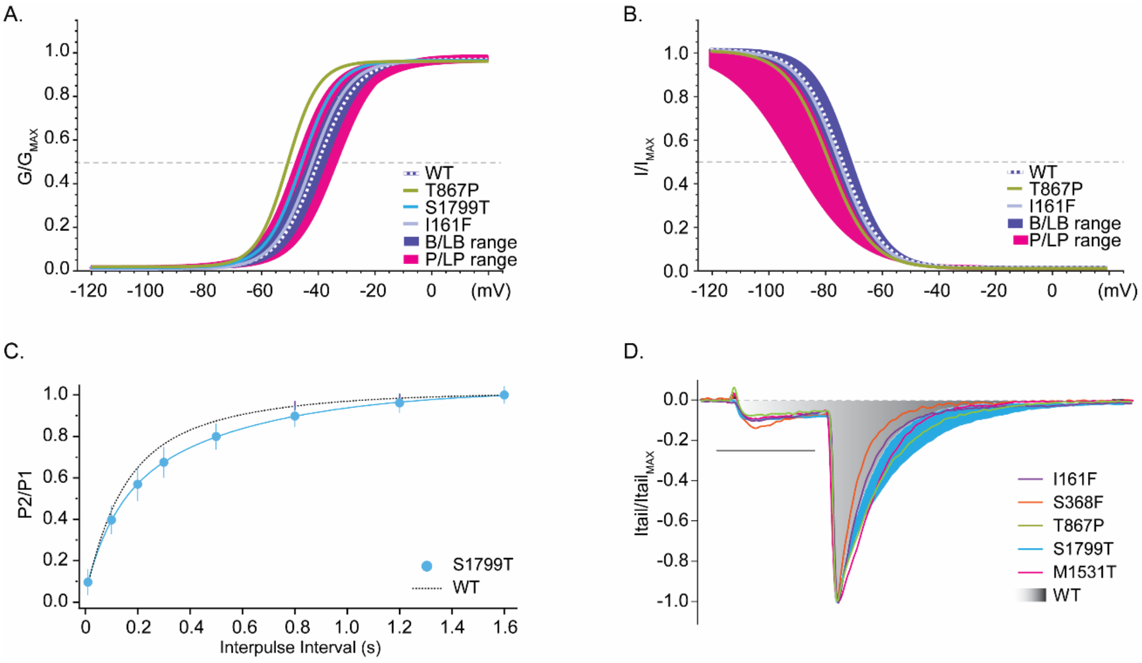


**Figure S3**. **A.** Normalized conductance–voltage relationships illustrating calibrated activation reference ranges, with benign/likely benign (B/LB) variants shown in magenta and pathogenic/likely pathogenic (P/LP) variants in blue. **B.** Normalized current-availability curves illustrating calibrated inactivation reference ranges, with B/LB variants shown in magenta and P/LP variants in blue. In panels **A** and **B**, Cav3.1 wild-type (WT) is shown as a dashed white line; variants of uncertain significance (VUS) with values overlapping the B/LB reference range are shown in yellow with white outlines, and VUS outside the B/LB range are shown in yellow with black outlines. **C.** Mean recovery from inactivation for Cav3.1-S1799T. Cav3.1-WT is shown as a dotted line for reference. **D.** Representative normalized current traces illustrating gating behaviour, scaled to maximal tail-current amplitude. The grey shaded gradient denotes the WT reference distribution. Scale bar: 10 ms.
