## Supplemental Figure 4 for "Calibrated high-throughput electrophysiology enables clinical interpretation of *CACNA1G* missense variants"

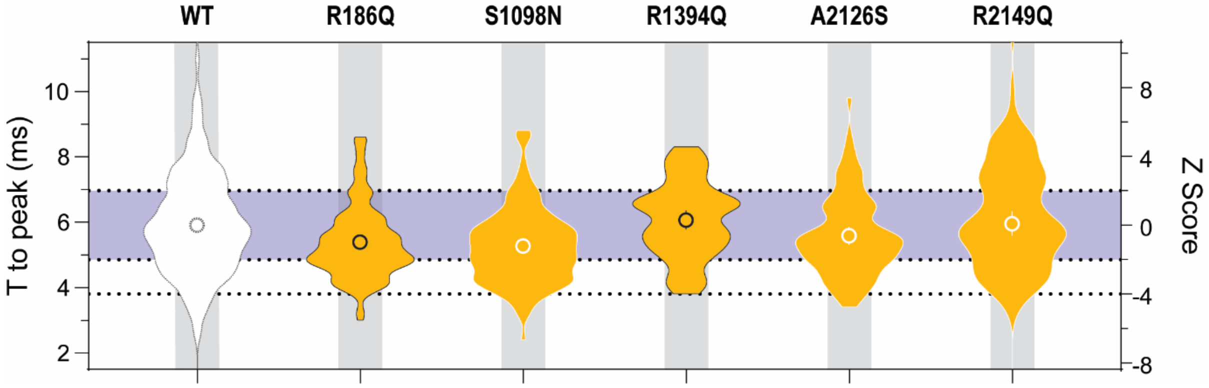


**Figure S4**. **Comparison of activation kinetics between Cav3.1-WT and VUS.** Mean ± 95% CI for time to peak are shown with raw values (left axis) and corresponding Z-scores (right axis). Normal functional range (mean ± 2 SD of benign values) is shaded in blue, and dotted lines indicate values beyond ±2. WT is shown in white and VUS in yellow. Variant-specific sample sizes are provided in Table S8.
